# MicrowellMicrofluidicsMiner (M³): Leverage Large Language Model Agents for Knowledge Mining of Microwell Microfluidics

**DOI:** 10.64898/2026.02.14.705953

**Authors:** Dinh-Nguyen Nguyen, Sadia Shakil, Raymond Kai-Yu Tong, Ngoc-Duy Dinh

**Author notes:** **Corresponding Author:** Corresponding, Address: BME office, Room 1120, 11/F, William M.W. Mong Engineering Building, or Room 208, Ho Sin Hang Engineering Building (SHB), The Chinese University of Hong Kong, Shatin, N.T., Hong Kong.

## Abstract

Microwell microfluidics has emerged as powerful platforms for high-precision biological and chemical investigations, bridging microscale fluid handling with compartmentalized reaction environments. Achieving robust and reproducible performance in such studies requires substantial effort to optimize microwell array design. This burden could be markedly alleviated by the availability of a curated database of microwell array parameters. Such a resource would enable the application of machine-learning models for performance prediction and automated design, leveraging knowledge accumulated from prior microfluidics research. However, constructing such a database entails a considerable investment of time and extensive manual curation, as microwell performance is governed by numerous critical design parameters that are reported inconsistently across a broad and largely unstructured body of literature. In this study, we introduce MicrowellMicrofluidicsMiner (M³), a framework that employs large language model (LLM) agents for autonomous knowledge extraction in microwell microfluidics. To evaluate its performance, we curate a ground-truth database and establish an LLM-driven assessment approach. Our results demonstrate that M³ achieves a peak accuracy of approximately 78%, representing more than a twofold improvement over the lowest observed accuracy (32%) obtained using a standalone LLM model (LLAMA 3.1). This study provides a foundational reference for researchers seeking to apply LLM agents to data-driven microfluidics research. The insights presented have the potential to substantially improve how scientists across microfluidics-related disciplines access, interpret, and leverage scientific information, thereby accelerating the development of innovative microfluidic devices and associated discoveries.

## Introduction

Microwell microfluidics is increasingly used for comprehensive single-cell investigations, spanning omics analyses and the assessment of cell–cell interactions^1–10^. This capability supports in-depth characterization of cellular heterogeneity, yielding enhanced understanding of diverse cell populations, complex tissue architectures, and disease heterogeneity. For instance, within the field of single-cell transcriptomics, a sealed microwell platform was employed to isolate mRNA from individual B cells using magnetic poly(dT) beads^11^. Subsequent emulsion PCR enabled the linkage and sequencing of paired heavy and light chains, thereby facilitating identification of B cell receptors (BCRs) at single-cell resolution. Moreover, the limitations of oil-based sealing were overcome through the development of Seq-Well, a microwell platform that employs a semi-permeable membrane for sealing^12^. This design is straightforward to implement and improves both transcript capture efficiency and platform portability. Regarding to single-cell proteomics, a strategy was developed to identify hybridoma cells that secrete antigen-specific antibodies by placing functionalized slides over microwell arrays^13^. After removing and imaging the slides, secretion profiles were mapped to individual microwells, allowing the isolation of viable, target-specific cells for subsequent clonal expansion. In addition, microbeads functionalized with DNA barcodes were incorporated into polydimethylsiloxane (PDMS) microwells to enable quantification of ten proteins at the single-macrophage level^14^. To achieve robust and reliable outcomes, these studies require substantial effort to optimise the design of microwell arrays. The burden of this labour-intensive step could be substantially reduced through the availability of a curated database of microwell array parameters. Such a database would facilitate the application of machine-learning models for performance prediction and automated design, drawing on insights from prior microfluidic research. **However**, ***the collection and extraction of this database demands a substantial investment of time and extensive manual curation, as microwell design depends on numerous key parameters (e.g., well dimensions, cell types and fabrication methods), each of which is reported inconsistently across a broad and largely unstructured literature*.**

Recent breakthroughs in generative artificial intelligence (AI) have led to the creation of advanced large language models (LLMs) trained on extensive datasets^15–18^. These models can analyze extensive text corpora, capture complex contextual relationships, and adapt to specialized domains^19–27^. This capability makes those LLMs particularly effective for automated extraction of data across many scientific fields. For example, Dagdelen *et al.*^28^ show that a fine-tuned LLM (GPT-3 or LLAMA-2) can jointly perform named-entity recognition and relation extraction to build materials-science databases. Similarly, Gupta *et al.*^29^ used GPT-3.5 and LLAMA-2 together with a MaterialsBERT named-entity-recognition model to extract over one million polymer-property records (e.g. tensile strength, glass transition temperature) from 681,000 polymer research articles. This pipeline turned free-form text into a structured polymer dataset, dramatically accelerating materials informatics efforts. Moreover, Murton *et al.*^30^ showed that an LLM could extract key trial parameters from clinical research reports almost as well as human reviewers. Likewise, Wiest *et al.*^31^ developed LLM-AIx, an end-to-end workflow where an LLM parses free-text pathology reports to pull out clinical entities. This pipeline enabled rapid, structured extraction of TNM staging data from hundreds of oncology reports without custom code. Additionally, Gougherty *et al.*^32^ reported that an LLM extracted emerging disease records from the literature over 50 folds faster than a human. The model pulled out discrete and categorical content with >90% accuracy. Also, Polak *et al.*^33^ introduced ChatExtract, a prompt-engineered pipeline that used a conversational LLM to locate relevant sentences, extract numerical data, and then verified each datum via follow-up questions.

However, a central obstacle limiting the reliability of LLM-based data extraction is the phenomenon of hallucination, whereby models generate plausible but incorrect or unsupported information^34^. Hallucination thus poses a serious risk when mining scientific data. To mitigate hallucination, retrieval-augmented generation (RAG) has emerged as a principled framework that explicitly grounds LLM outputs in external evidence^35^. In a RAG system, the LLM is paired with a retrieval component that fetches relevant documents or data from an external corpus, and the model’s generation is conditioned on this retrieved evidence^35^. Beyond RAG, to further enhance robustness, researchers have increasingly turned to large language model agents, which extend LLMs from passive text generators into autonomous systems capable of multi-step reasoning, executing actions and tool use^36–39^. For example, the SciAgents framework^40^ leverages multiple specialized LLM agents to generate original research ideas, develop experimental plans, and perform refinement of scientific hypotheses. Within the chemical research, ChemCrow^41^ represents an LLM-driven chemistry agent developed to advance scientific discovery by bridging experimental and computational approaches. The system combines an LLM with a set of 18 chemistry tools developed by domain experts, including molecular property predictors, reaction planning systems, and chemical databases, thereby enabling the autonomous planning and execution of chemical syntheses. In biology domain, GeneAgent^42^ leverages a self-refinement circle to uncover associations between genes derived from biomedical datasets, thereby increasing the reliability of its results through validation against established gene sets. ***While LLM agents are increasingly applied in other areas, their adoption for automated data extraction in microwell microfluidics is limited, emphasizing a significant opportunity for innovation*.**

In this study, we propose MicrowellMicrofluidicsMiner (M³), an LLM agent-driven framework for autonomous knowledge extraction in microwell microfluidics. The framework integrates three core components: a retrieval-augmented generation (RAG) module, a mixture-of-agents (MoA) ensemble^43^, and an LLM extractor. The RAG module ensures retrieval of relevant information to reduce hallucinations, whereas the MoA ensemble incorporates diverse model capabilities and learned linguistic patterns. This ensemble design introduces reasoning diversity, reduces biases inherent to individual models, and offers redundancy for robust performance. The LLM extractor evaluates the quality of responses through prompt assessment. **We introduce an LLM-driven assessment framework that alleviates human workload across high-throughput settings such as iterative model development, large-scale data extraction, and ongoing system monitoring**. This work offers a foundational contribution for researchers aiming to apply LLMs to data-driven microfluidics research. By improving access to and utilization of scientific knowledge across related disciplines, the findings have the potential to accelerate the development of novel microfluidic devices and scientific discovery.

Add: agent evaluator

## 2. Methodology

### 2.1 Selection of models

To maintain stringent scientific standards, the study relied solely on open-source models, incorporating five foundational LLMs into the M³ framework to ensure transparency, complete accessibility to model architectures and weights, and full reproducibility. By eliminating reliance on commercial APIs and their associated costs, the workflow remains both scalable and cost-effective, thereby supporting extended experimentation and promoting broad community adoption while preserving core principles of openness and reproducible research. The LLMs incorporated in this study are enumerated in detail in Table 1.

**Table 1.** Applied LLMs.

| Model name | Size | Provider | Open | Source |
| --- | --- | --- | --- | --- |
| GEMMA2 | 9B | Google | Yes | <sup>44</sup> |
| LLAMA3.1 | 8B | Meta | Yes | <sup>17</sup> |
| MISTRAL | 7B | Mistral AI | Yes | <sup>45</sup> |
| DEEPSEEK-R1 | 8B | DeepSeek | Yes | <sup>18</sup> |
| QWEN3 | 14B | Alibaba | Yes | <sup>46</sup> |

### 2.2 MicrowellMicrofluidicsMiner (M³) framework

The process of M³ framework begins with a user-defined query specifying the parameters of interest, which is encoded and routed through an RAG module that interfaces with a domain-specific vector database, as shown in Fig. 1(a). This module identifies semantically relevant documents based on high-dimensional embeddings and returns text segments that potentially contain the requested information. To improve precision, the retrieved passages undergo a reranking stage, which prioritizes sources with the strongest evidence for the queried parameter. The highest-ranking passages are then summarized to remove redundancy, harmonize terminology, and produce a concise, context-optimized representation that can be reliably ingested by downstream models. This refined context is subsequently evaluated by a mixture of independent LLM agents, including LLAMA3.1, MISTRAL, DEEPSEEK-R1, and GEMMA2. Each of which produces an answer candidate informed by distinct architectural biases, reasoning strengths, and linguistic capabilities. The resulting model-diverse answer set is forwarded to a QWEN3-based extractor, which performs cross-model arbitration by selecting or synthesizing the response that most closely aligns with the retrieved evidence and the original information need. In detail, the QWEN3-based extractor uses the prompt evaluation to assess the quality of responses generated by four LLMs to a given question, using a specified retrieved context, as illustrated in Fig. 1(b). First, the extractor applies a majority voting rule, automatically choosing any response shared by at least two LLMs. If no majority exists, the extractor uses a tie-breaking rule that favours responses that are more strongly supported by the provided context and articulated with superior clarity, conciseness, and structural coherence. If all four responses differ, the extractor invokes a fallback mechanism, selecting the response that is most contextually relevant and factually accurate, applying the tie-breaking criteria if needed. The final output must consist solely of the selected response. This arbitration step substantially reduces hallucination risk and enhances factual grounding by leveraging both consensus signals and cross-model complementarity. The details of the prompt used for the QWEN3-based extractor is provided in the supplementary file (Table S1).

**Fig. 1.**
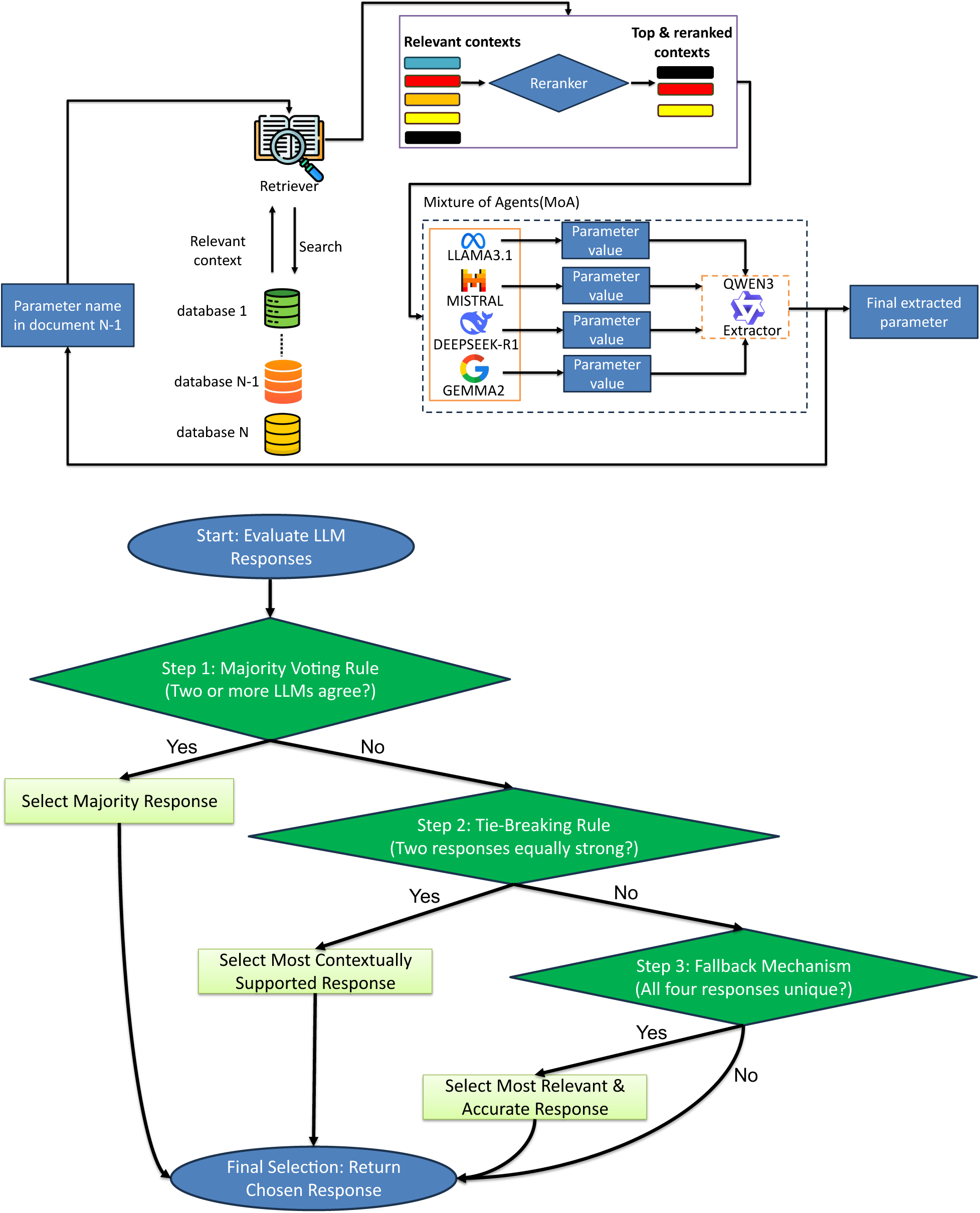
(a) MicrowellMicrofluidicsMiner (M³) framework. (b) Flowchart of prompt used for the QWEN3-based extractor

### 2.3 Curated ground-truth construction

In this study, we compiled a ground-truth dataset informed by a previous investigation^47^. The dataset was derived from 39 peer-reviewed publications and captures six key parameters relevant to microwell experiments: microwell material, microwell shape, fabrication method, bioapplication, cell type, and microwell dimension. In total, the dataset contains 234 entries, which we used to evaluate the performance of the M³ framework. The details of the dataset is provided in the supplementary file (Table S2).

### 2.4 Database construction for RAG

The flowchart outlines a structured pipeline for transforming unprocessed scientific documents into a searchable knowledge substrate suitable for RAG systems, as illustrated in Fig. 2. The process begins with a *data source*, such as PDF files, which frequently contain complex layouts, mixed text–figure regions, and diverse formatting conventions. These challenges require a dedicated *information extraction* stage, which converts the heterogeneous PDF content into clean, logically ordered text. The resulting text is then passed to a *chunking* module, which segments the content into semantically coherent units of manageable length. Each chunk or unit is subsequently fed into an *embedding model*, typically a transformer-based encoder, that maps textual segments into dense vector representations capturing their semantic relationships. The resulting *embeddings* are stored in a *vector database*.

**Fig. 2.**
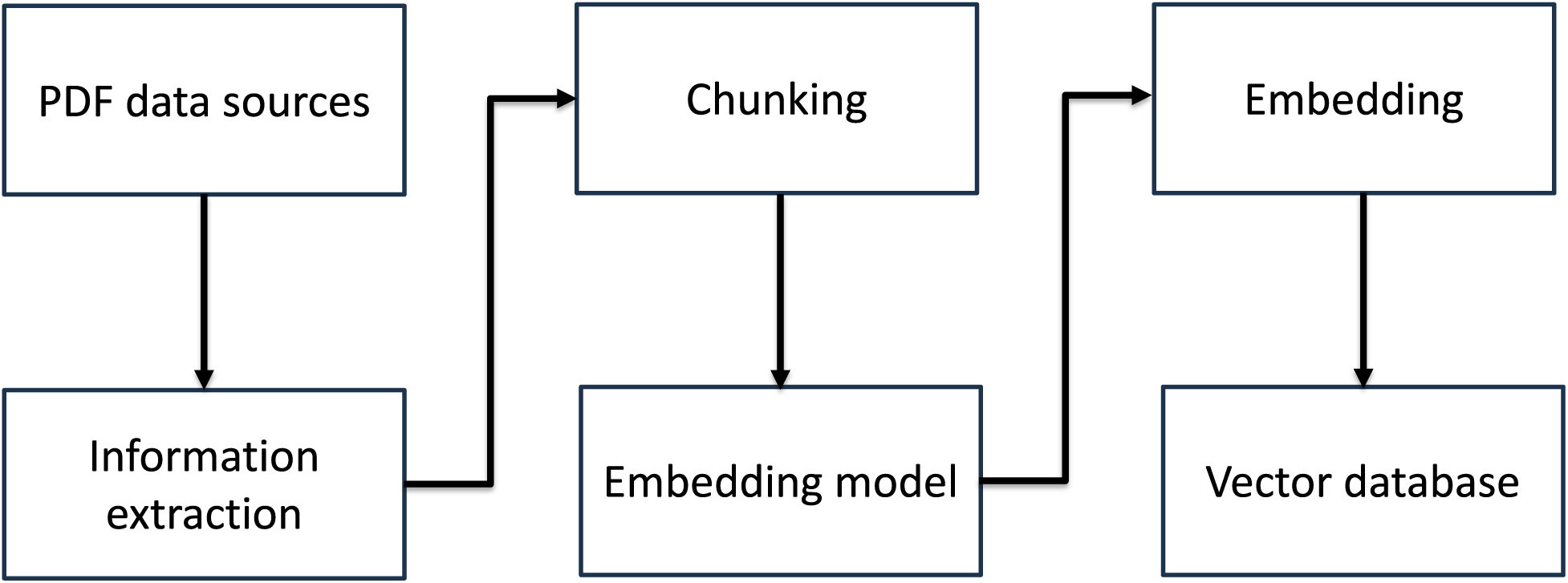
Flowchart of database construction for RAG

### 2.5 LLM as a judge

The schematic outlines an evaluation pipeline in which the M³ framework is tested against a ground-truth dataset using an LLM judge based on QWEN3 to provide accuracy scores, as shown in Fig. 3. The process begins with a predefined set of questions. Each question is linked to a validated ground-truth answer, typically derived from expert review or high-quality reference datasets, which anchors the evaluation in a reliable standard. The same question is then fed into M³ framework. Once generated, the M³ output and its corresponding ground truth are simultaneously provided to an independent LLM agent serving as a judge. This judge performs a semantic comparison that goes beyond lexical matching, evaluating the extent to which the extracted answer captures the factual, contextual, and logical content of the ground truth. When the response is largely relevant, coherent, and substantively accurate, the judge assigns a correspondingly high score, while ensuring that partially correct answers, particularly those capturing the essential intent but lacking detail, still receive a fair, above-median assessment. The judge also provides constructive feedback that identifies key strengths and gently notes any areas where clarity, completeness, or precision could be improved. To maintain consistency, the evaluation output follows a standardized format: *“Feedback: {{feedback}} SCORE: {{0–100}}”.* The details of the prompt used for the judge are provided in the supplementary file (Table S3). We further conducted a direct comparison between the accuracy scores assigned by the LLM-based evaluation agent and those provided by a microfluidics expert in assessing the extraction results generated by the M³ framework. Detailed analyses of this comparison are presented in Section 3.

**Fig. 3.**
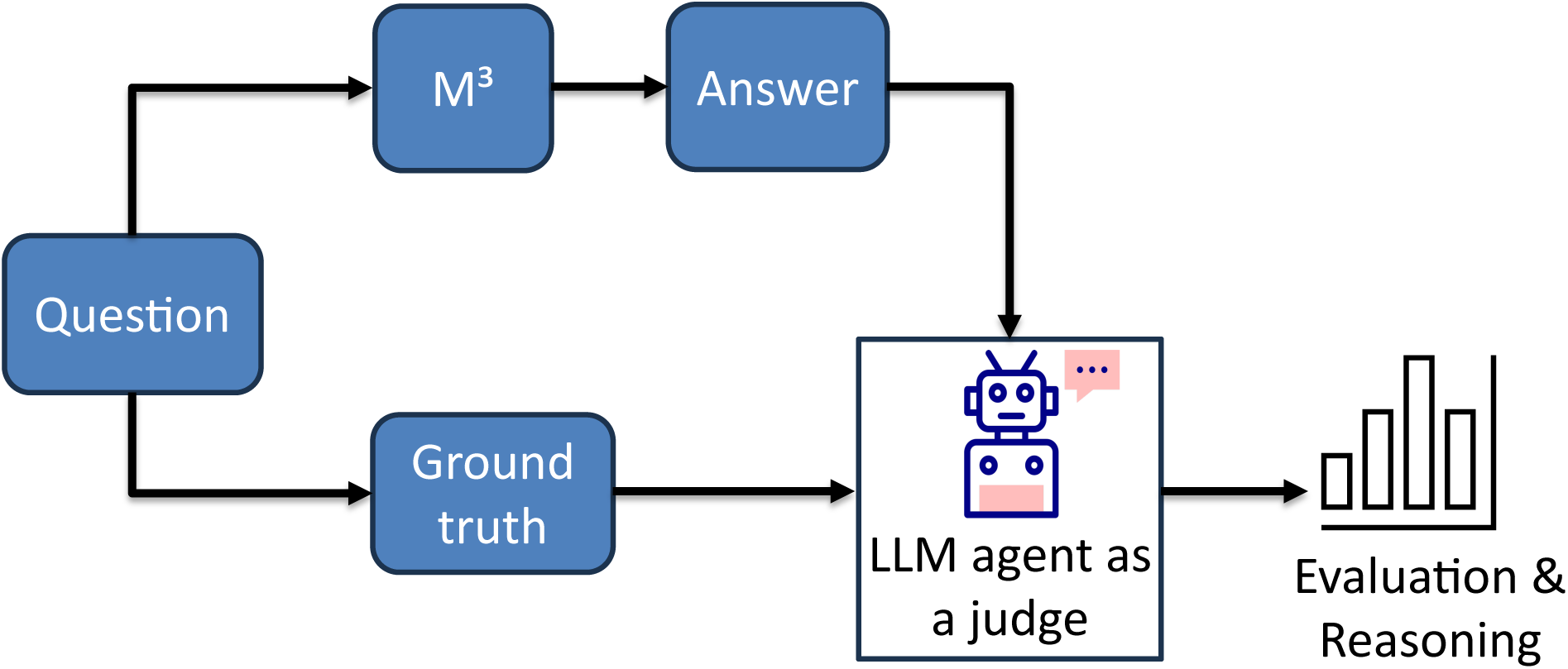
LLM as a judge

### 2.6 Investigation of the impact of embedding models on the accuracy of the M³ framework

The selection of an embedding model plays a critical role in retrieval performance, as it determines how effectively semantic relationships between user queries and document segments are encoded. In this work, we systematically evaluated the impact of embedding model choice on the extraction accuracy of the M³ framework for microwell dimension parameter. Performance was assessed using seven open-source embedding models: *intfloat-e5-large-v2*^48^, *all-MiniLM-L6-v2*, *all-mpnet-base-v2*, *pritamdeka-BioBERT-mnli-snli-scinli-scitail-mednli-stsb*^19^, *BAAI-bge-base-en-v1.5*, *msmarco-distilbert-dot-v5*, and *sentence-transformers-stsb-roberta-base-v2*^49^. All models are publicly available via the Hugging Face repository(https://huggingface.co).

## 3. Results & discussion

A representative example of parameter extraction using the M³ framework, showcasing how the system synthesizes retrieved evidence and multi-LLM reasoning to identify the experimental dimensions of microwells reported in a published study, is illustrated in Fig. 4. The retrieved text explicitly states a microwell depth of 120 µm, a diameter of 300 µm, and an inter-well spacing of 1 mm, and these values are consistently recovered across all four contributing LLMs, GEMMA2, Mistral, LLAMA3.1, and DeepSeek-R1, despite minor stylistic or inferential differences. Notably, Qwen3-Extractor, serving as the arbitration model, converges on the complete and numerically accurate parameter set, mirroring the ground-truth values with no deviation. The LLM-judge’s evaluation assigns a score of 95, acknowledging near-perfect alignment with the ground truth, while expert review yields a score of 100, confirming the fidelity of the extracted parameters. Collectively, these results highlight the robustness of the M³ framework in extracting experimental conditions and underscore its potential utility for large-scale, automated synthesis of methodological metadata across the scientific literature.

**Fig. 4.**
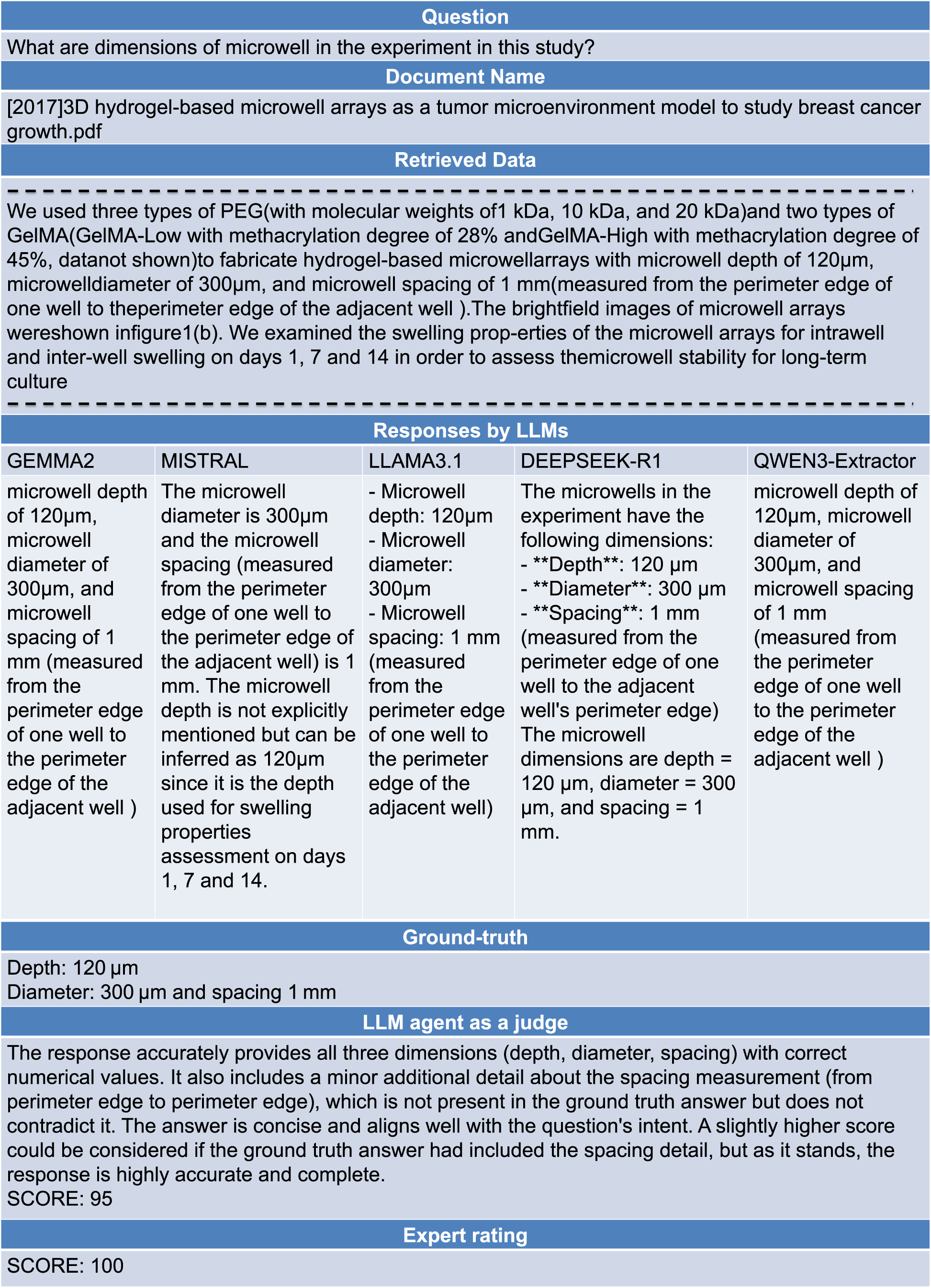
An illustration example of parameter extraction using M³

The summary summarizes the extraction accuracy of the M³ framework across six key microwell parameters and simultaneously underscores several important advantages of the approach, illustrated in Fig. 5. Accuracy values span from 60% to 78%, with bioapplication achieving the highest accuracy of around 78%, outperforming the lowest-scoring parameter microwell dimension of about 60%. High performance for microwell material of roughly 76% and fabrication technique of roughly 72% demonstrates that the system reliably identifies parameters grounded in standardized terminology and well-defined methodological phrasing. Meanwhile, moderate but consistent accuracies for microwell shape of approximately 64% and cell name of close to 69% reflect the model’s capacity to extract semantically diverse descriptors despite substantial lexical variability across the literature. Importantly, the results reveal several clear advantages of the M³. First, despite heterogeneity in parameter types, the framework maintains accuracies consistently above 60%, underscoring its robustness across both engineering-focused and biologically oriented descriptors. Second, its strong performance on both qualitative (e.g., shape, application) and quantitative or process-driven parameters (e.g., dimensions, fabrication technique) shows that the method is not restricted to narrow descriptor classes, positioning it well for broad-spectrum literature mining in microfluidics. Third, the high accuracy for material, application, and fabrication parameters suggests that the architecture effectively integrates contextual cues and domain-specific semantics, enabling meaningful extraction without extensive manual feature engineering. Fourth, because it generalizes well across disparate categories, the M³ is immediately scalable, supporting high-throughput analysis of hundreds to thousands of publications, an advantage that traditional manual curation or rule-based systems cannot match. In a broader scientific context, these strengths demonstrate that the M³ meaningfully advances automated knowledge extraction in microfluidics research. Its combination of robustness, domain adaptability, and parameter diversity enables researchers to rapidly construct structured datasets, explore design trends, and conduct systematic meta-analyses with far greater efficiency than conventional methods, ultimately accelerating the pace of data-driven innovation in microwell microfluidic systems.

**Fig. 5.**
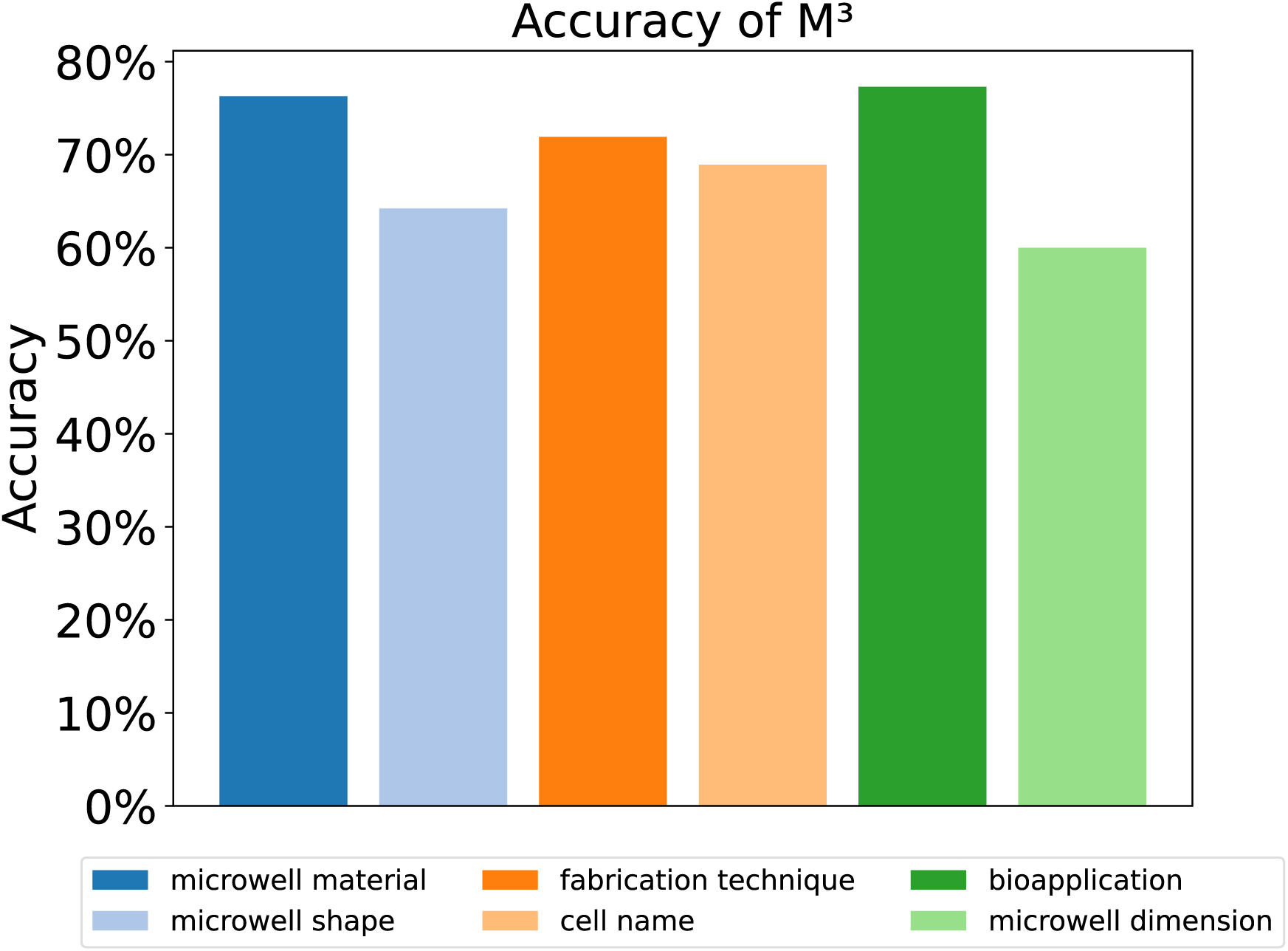
Extraction accuracy of the M³ framework

A comparative evaluation of parameter extraction accuracy for fabrication technique parameter across five state-of-the-art LLMs, DEEPSEEK-R1, GEMMA2, LLAMA3.1, MISTRAL, and QWEN-3, benchmarked against the M³, is shown in Fig. 6. Whereas standalone LLM accuracies range only from 32% to 55%, M³ achieves approximately 72% exceeding the weakest by more than 2-fold. These results underscore the persistent difficulty that general-purpose LLMs face in extracting fine-grained parameters from complex scientific manuscripts. Collectively, these findings position the M³ as a significantly more reliable, interpretable, and scalable solution for large-scale extraction of parameters from scientific literature in microwell microfluidics.

**Fig. 6.**
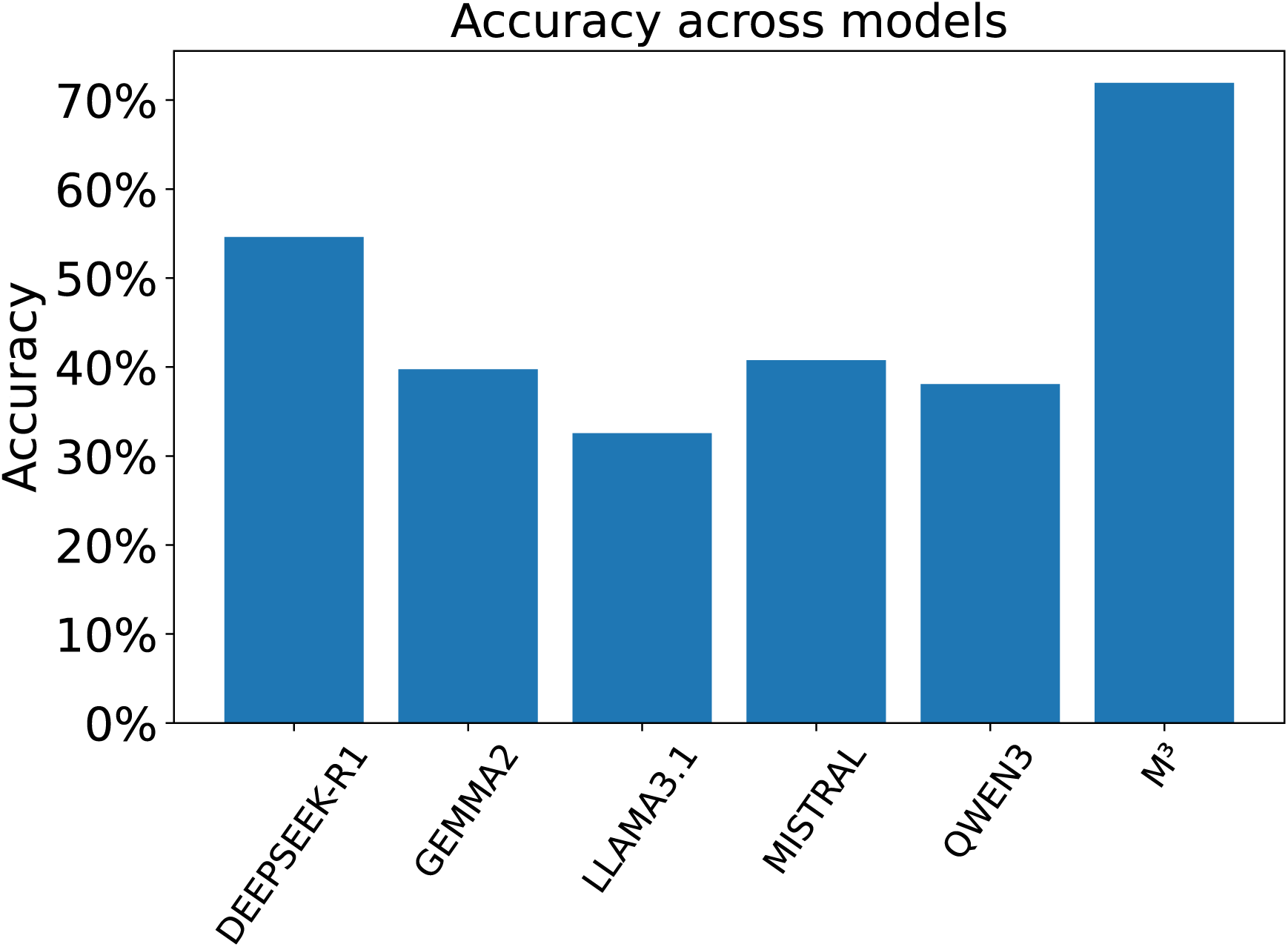
Extraction accuracy of M³ against standalone LLMs

A comparative analysis of microwell dimension parameter extraction accuracy across seven sentence-embedding models within the M³ framework is shown in Fig. 7. Accuracy varied substantially among models, ranging from approximately 20% for *all-MiniLM-L6-v2* to nearly 63% for *all-mpnet-base-v2*, representing a more than threefold increase in performance. Several models including *msmarco-distilbert-dot-v5* (around 60%) and *intfloat-e5-large-v2* (about 60%) exhibited similarly high accuracies, whereas domain-oriented biomedical encoders such as *pritamdeka-BioBERT-mnli* (approximately 55%) and *sentence-transformers-stsb-roberta-base-v2* (close to 52%) showed moderate performance. **The highest accuracy of *all-mpnet-base-v2* in the parameter-extraction task arises from a combination of architectural design, pretraining strategy, and representational capacity that aligns particularly well with the semantic demands of scientific text and RAG-based workflows. At the architectural level, *all-mpnet-base-v2* integrates masked language modeling with permuted token prediction, enabling the model to learn bidirectional context while simultaneously preserving word-order dependencies. This hybrid objective allows the embedding space to encode both fine-grained local semantics for example relationships between parameters and their units and long-range contextual structure such as conditions, comparisons, and methodological dependencies spread across sentences. Critically, *all-mpnet-base-v2* is trained on large-scale corpora optimized for semantic similarity and sentence-level reasoning. This approach enhances its ability to generalize across diverse writing styles, experimental formats, and numerical reporting conventions found in research papers. Equally important is the high representational capacity of *all-mpnet-base-v2*. With deeper transformer layers and higher-dimensional embeddings than compact models such as *all-MiniLM-L6-v2*, *all-mpnet-base-v2* can encode subtle semantic distinctions between closely related experimental descriptions. This directly improves retrieval discrimination in RAG pipelines**. Collectively, these results suggest that general-purpose, high-capacity embedding models outperform lightweight or domain-tuned counterparts for this task, highlighting that effective parameter mining from research literature may rely more on semantic expressiveness than on domain specialization. They also underscore the importance of model selection for scientific information extraction.

**Fig. 7.**
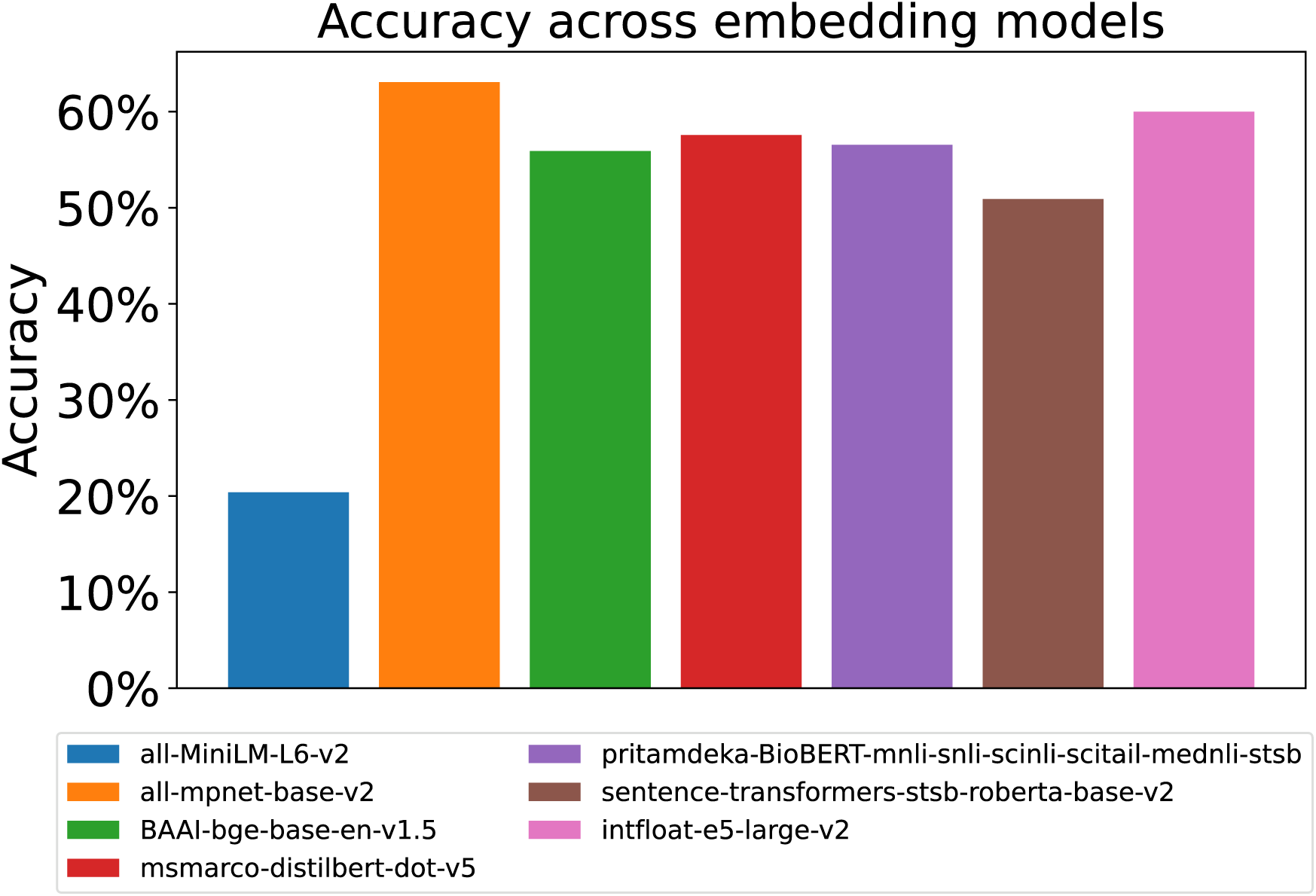
Extraction accuracy across sentence-embedding models

A direct comparison between accuracy scores of fabrication technique parameters assigned by the LLM-based judging agent powered by QWEN3 and those determined by a microfluidics expert for the grading of extraction results generated by the M³ framework, is shown in Fig. 8. The resulting regression line demonstrates a near-ideal linear relationship, yielding an R² value of 0.98, which indicates that 98% of the variance in expert judgments is captured by the LLM’s evaluations. Such a high coefficient of determination demonstrates that the LLM agent closely mirrors expert decision-making across the full accuracy range. Taken together, these results offer compelling evidence that LLM-based evaluation is technically feasible and operationally advantageous for large-scale grading tasks that would otherwise require substantial expert labor. The strong alignment with expert scoring implies that the LLM agent can serve as a reliable first-pass evaluator, reducing the human workload in high-throughput assessment scenarios such as iterative model development, large-batch information extraction, and continuous system monitoring.

**Fig. 8.**
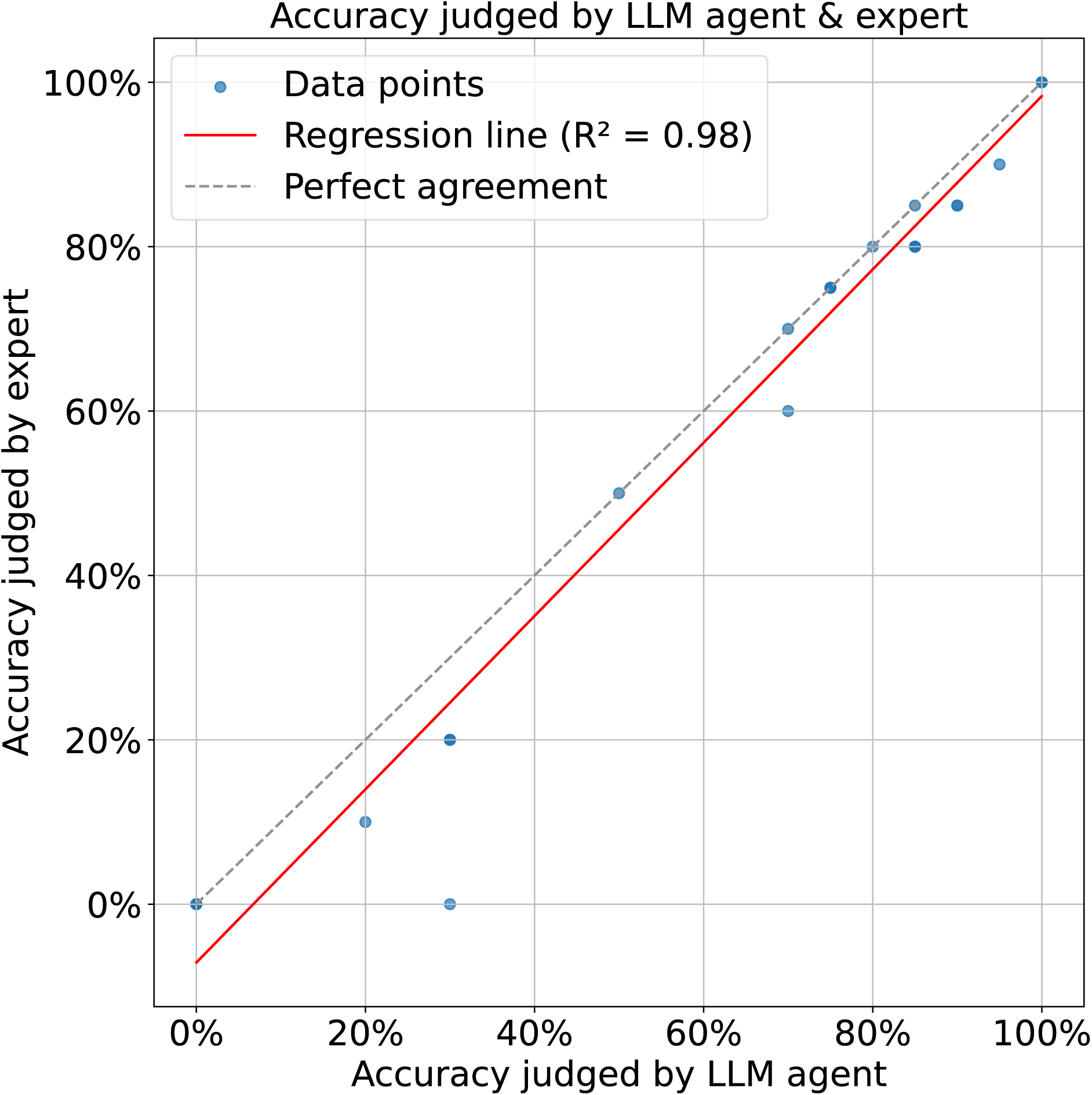
Accuracy comparison of an LLM-based judging agent and a microfluidics expert

## 4. Conclusion and Outlook

In this study, we present M³, a LLM agent-driven framework designed for autonomous knowledge extraction in microwell microfluidics. M³ addresses a fundamental challenge in the field: although a substantial body of microwell microfluidics knowledge is embedded in unstructured natural language, structured representations are essential for enabling systematic analysis, performance prediction, design automation, and downstream discoveries. The framework comprises three core components: RAG module, MoA ensemble, and an LLM-based extractor. Experimental results demonstrate that M³ achieves a peak extraction accuracy of approximately 78%, corresponding to more than a twofold improvement over the lowest observed accuracy (32%) obtained using a standalone LLM (LLAMA3.1). We further conduct a systematic empirical analysis to evaluate the influence of different embedding models on extraction performance. Among the models assessed, *all-mpnet-base-v2* yields the highest accuracy. In addition, we directly compare accuracy scores produced by an LLM-based judging agent with those assigned by a microfluidics expert when evaluating extraction outputs generated by the M³ framework. Linear regression analysis reveals a near-ideal correspondence between the two, with an R² value of 0.98, indicating that 98% of the variance in expert assessments is explained by the LLM’s evaluations. This high coefficient of determination demonstrates that the LLM-based judging agent closely approximates expert decision-making across the full accuracy range. This study establishes a foundational resource for researchers seeking to leverage LLMs for data-driven investigations in microwell microfluidics and highlights the potential of LLM agents to transform how scientific knowledge is accessed, structured, and applied across microfluidics research, thereby accelerating the development of novel devices and discoveries addressing critical societal challenges.

Microfluidics, encompassing channel-based^50–52^, droplet-based^53–59^, and microwell-based platforms, is a data-intensive field that nevertheless faces persistent challenges in data curation and standardization. Microfluidic devices and experiments are typically reported with extensive quantitative detail, including geometric parameters (such as channel and microwell dimensions), operational conditions (for example, flow rates and fluid properties), and experimental outcomes (such as droplet size and generation rate). However, this information remains fragmented across the literature, limiting its reuse for large-scale analysis.

Initial efforts to compile microfluidics datasets for machine learning applications have begun to address this gap. For example, Lashkaripour *et al.*^60,61^ curated a droplet microfluidics dataset and trained machine learning models to predict device designs capable of producing target droplet sizes. LLM-based approaches offer a powerful means to further streamline this process by automatically parsing published methods and extracting relevant numerical values together with their associated units. In this context, Nguyen *et al.*^62^ advanced the field with μ-Fluidic-LLMs, which convert tabulated experimental records into natural-language descriptions and subsequently leverage pre-trained LLMs to encode and extract informative features. This hybrid strategy produced substantial performance gains, with the inclusion of LLM-derived contextual information significantly reducing prediction errors for droplet diameter and generation rate relative to previous models. These early efforts show that LLMs can bridge between human language descriptions and structured microfluidic data. By automating the laborious process of literature review and data entry, such LLM agents have the potential to accelerate microfluidic design and discovery^63^.

## Data availability

The datasets and code for the analyses and figure generations in this work are publicly available on GitHub at url: https://github.com/duydinhlab/MicrowellMicrofluidicsMiner

## Author Contributions

Conceptualization – N-D.D.; Methodology – D-N.N., N-D.D.; Investigation – D-N.N.; Data curation – D-N.N.; Writing – original draft – D-N.N.; Writing – review & editing – D-N.N., N-D.D.; Supervision – S.S., R.K-Y.T., N-D.D; Funding acquisition – R.K-Y.T., N-D.D.

## Acknowledgements

We gratefully acknowledge the funding provided by the Research Grant Council of Hong Kong, General Research Fund (Ref No. 14211223).

## Table of Content

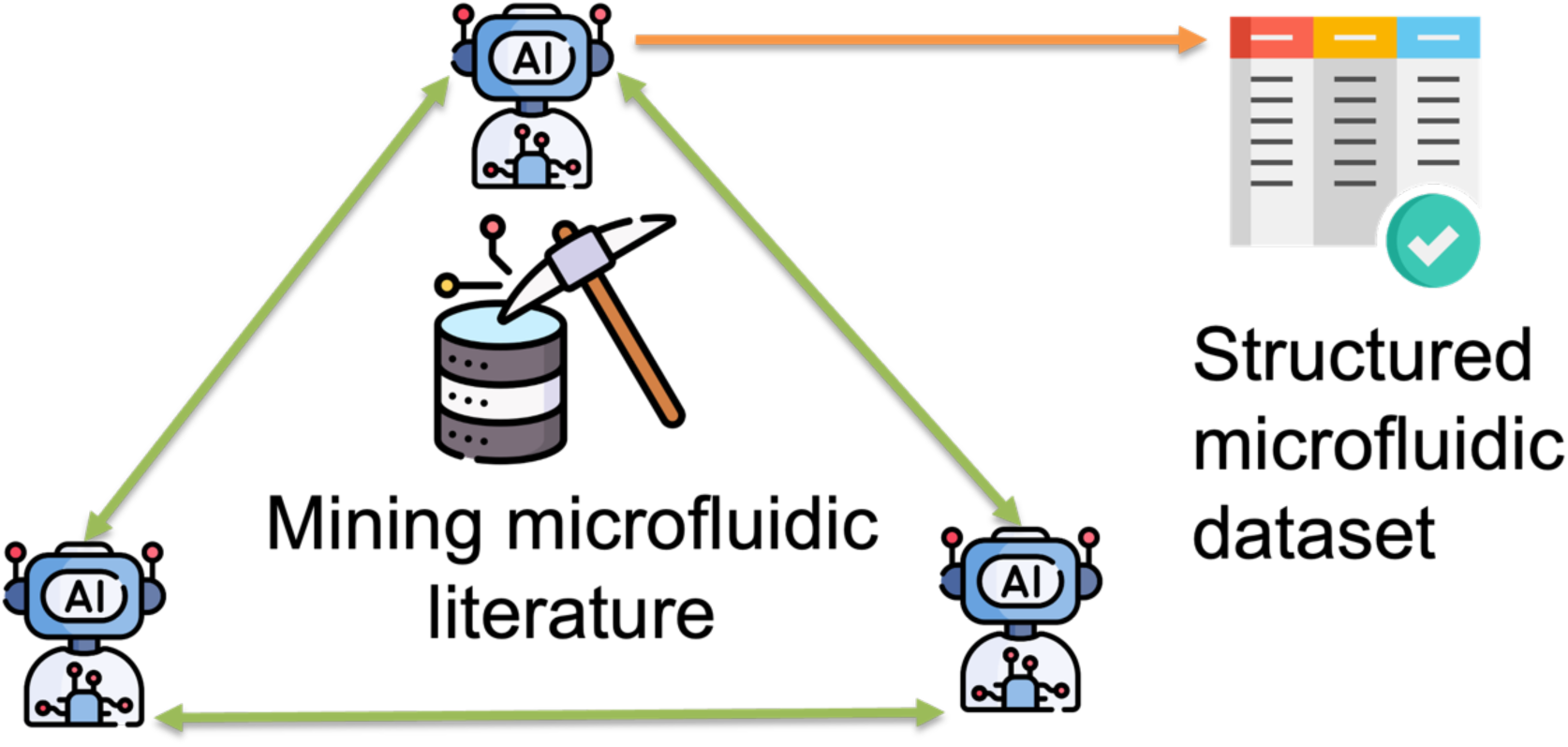

## Supplementary File

**Table S1.**
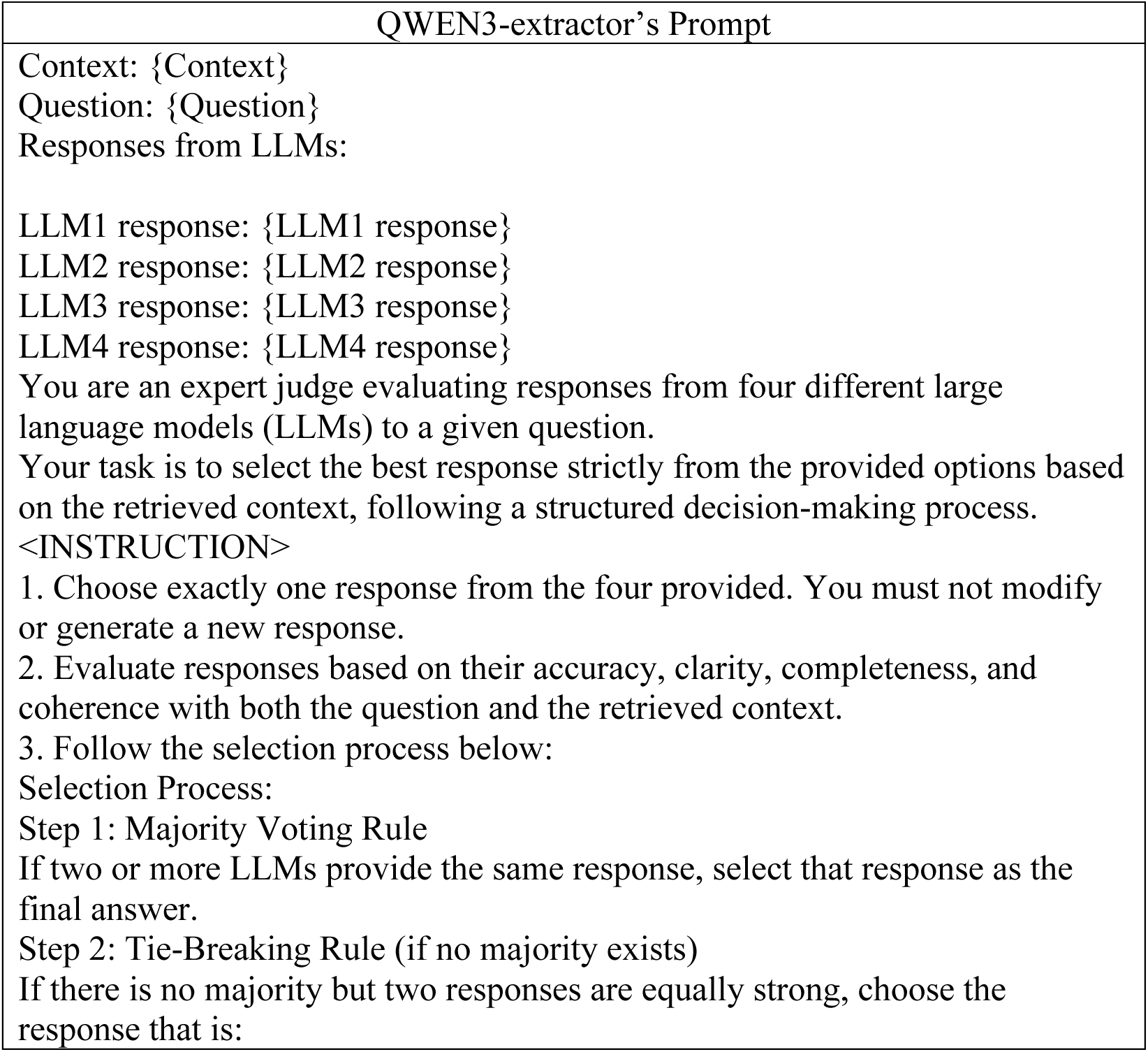
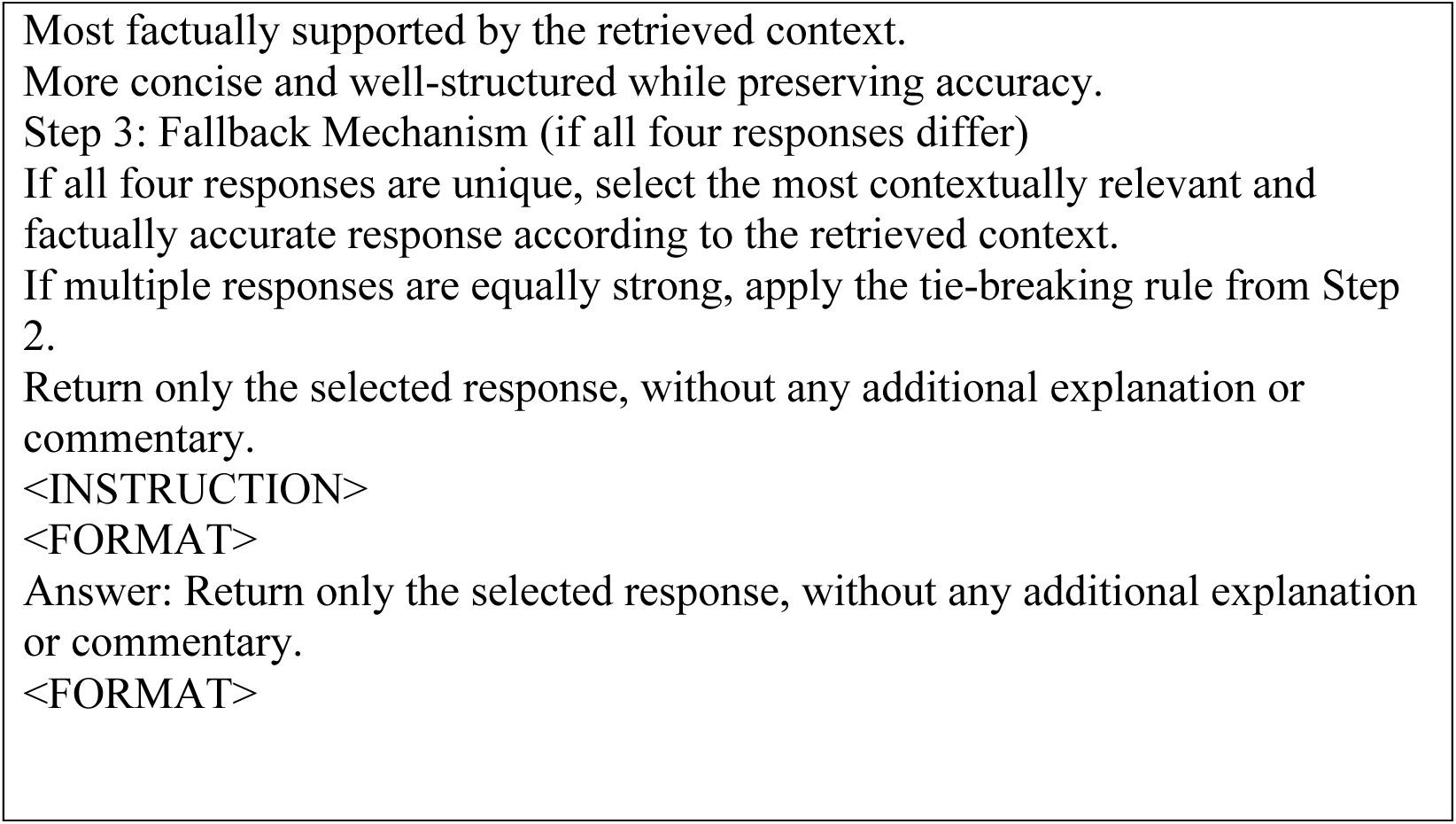
QWEN3-extractor’s Prompt.

**Table S2.**
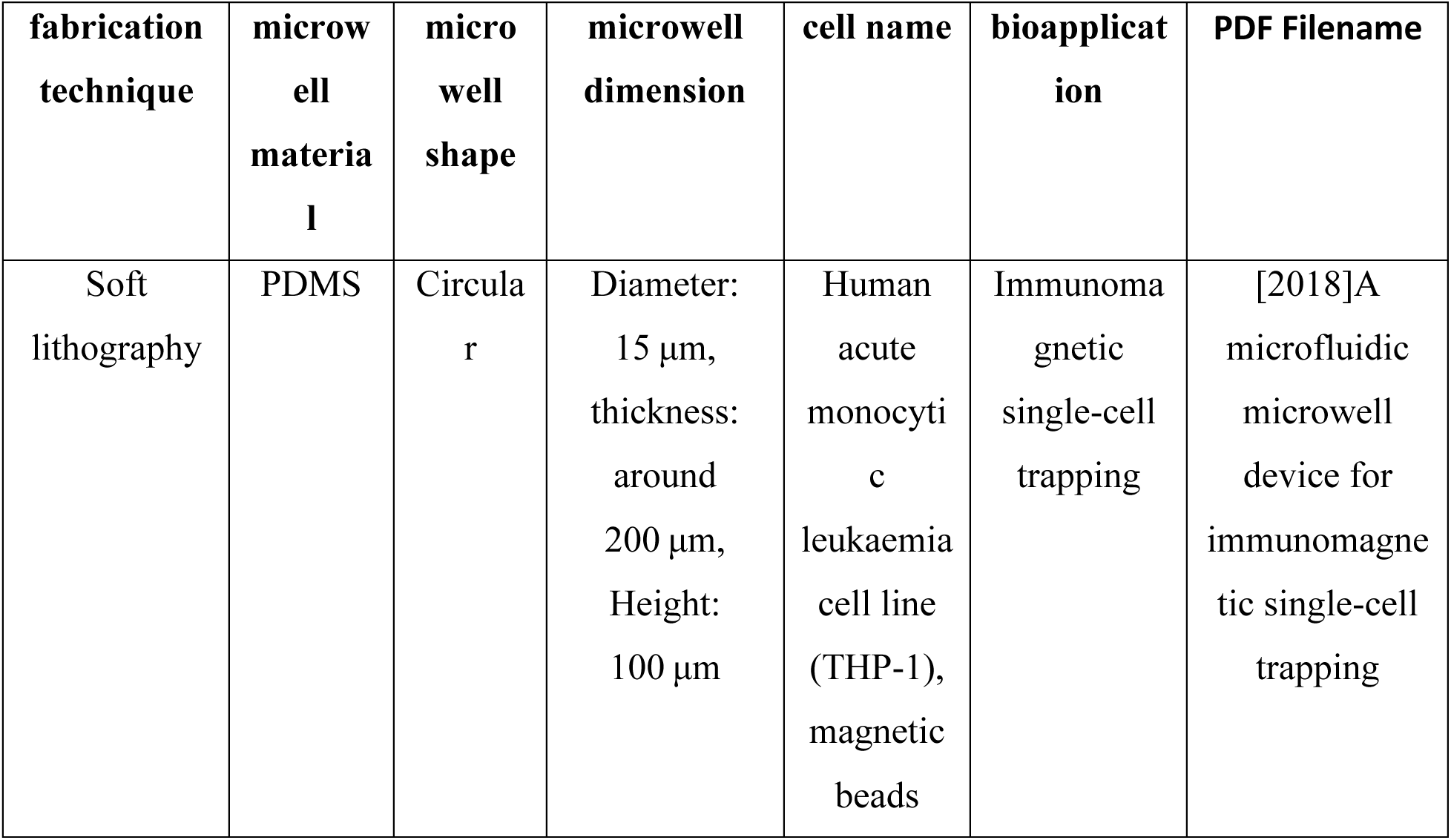
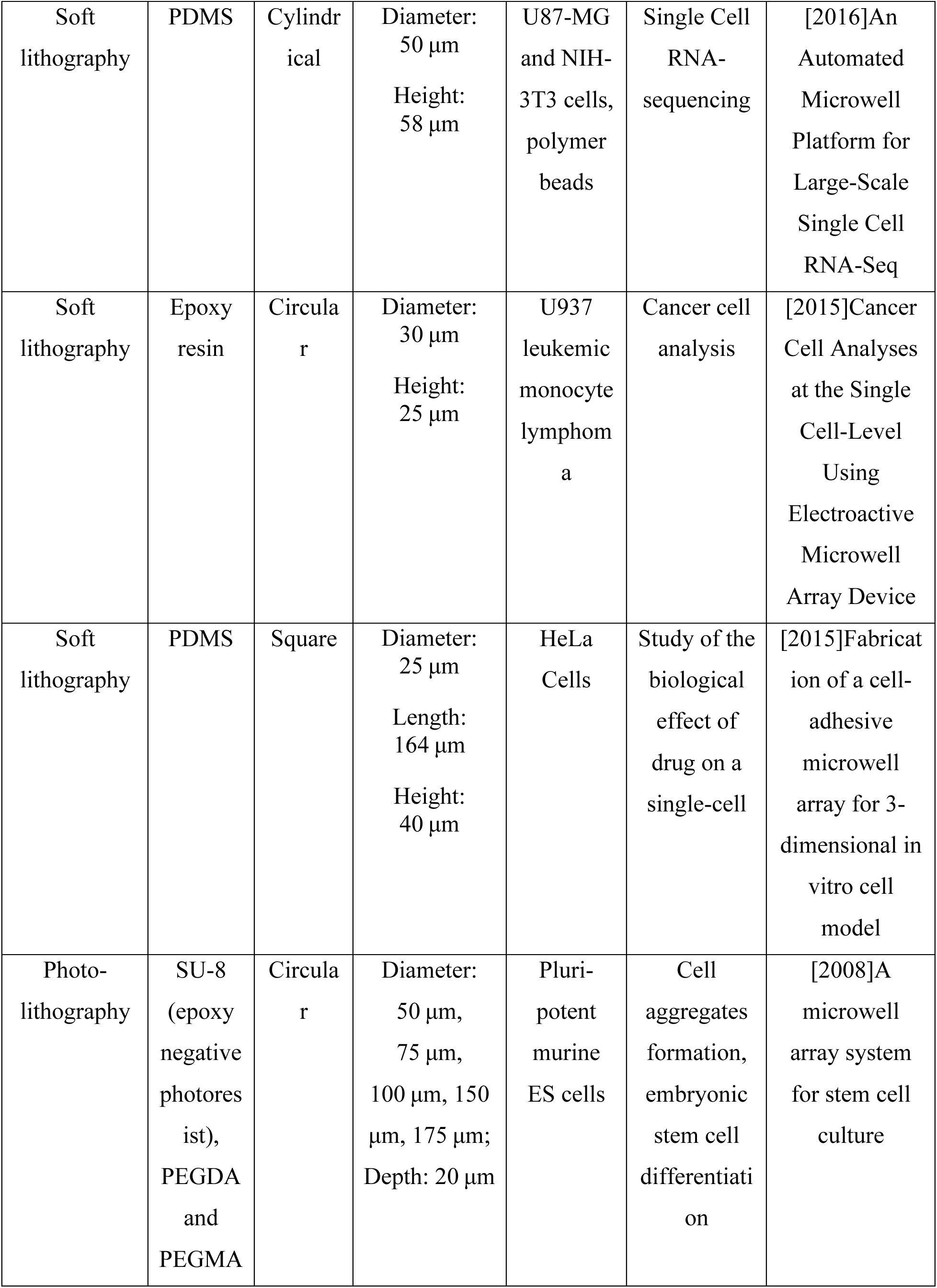
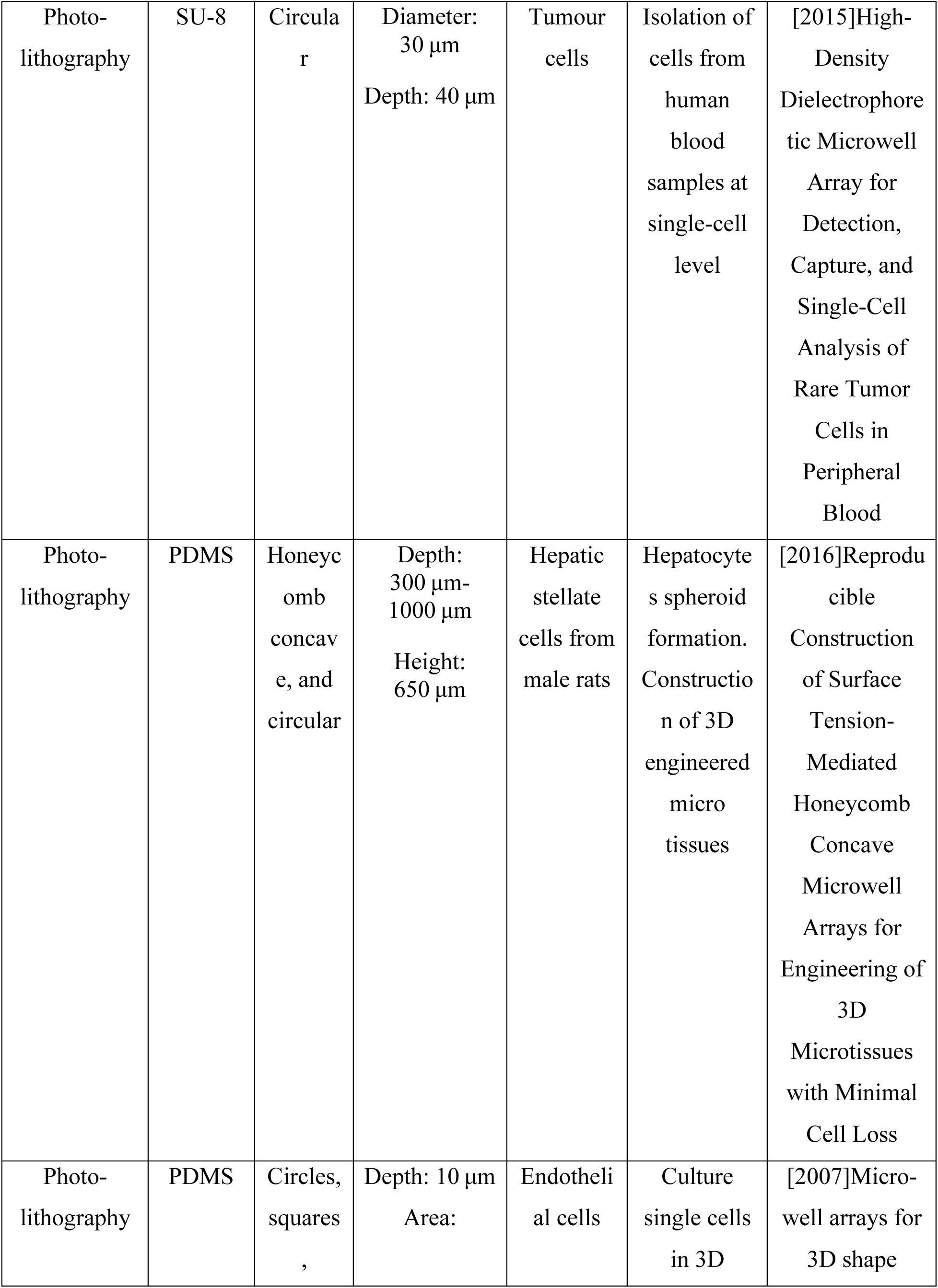
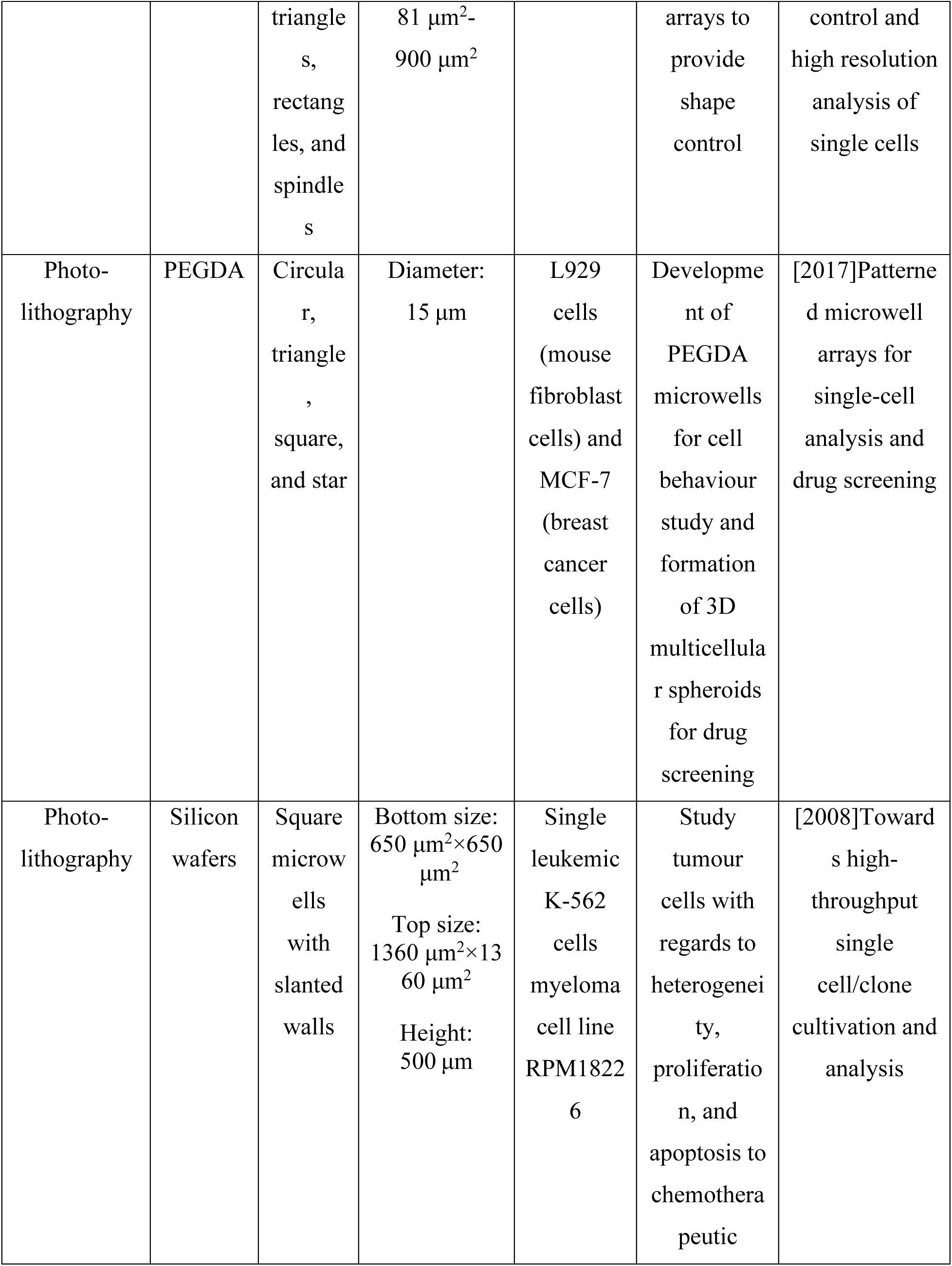
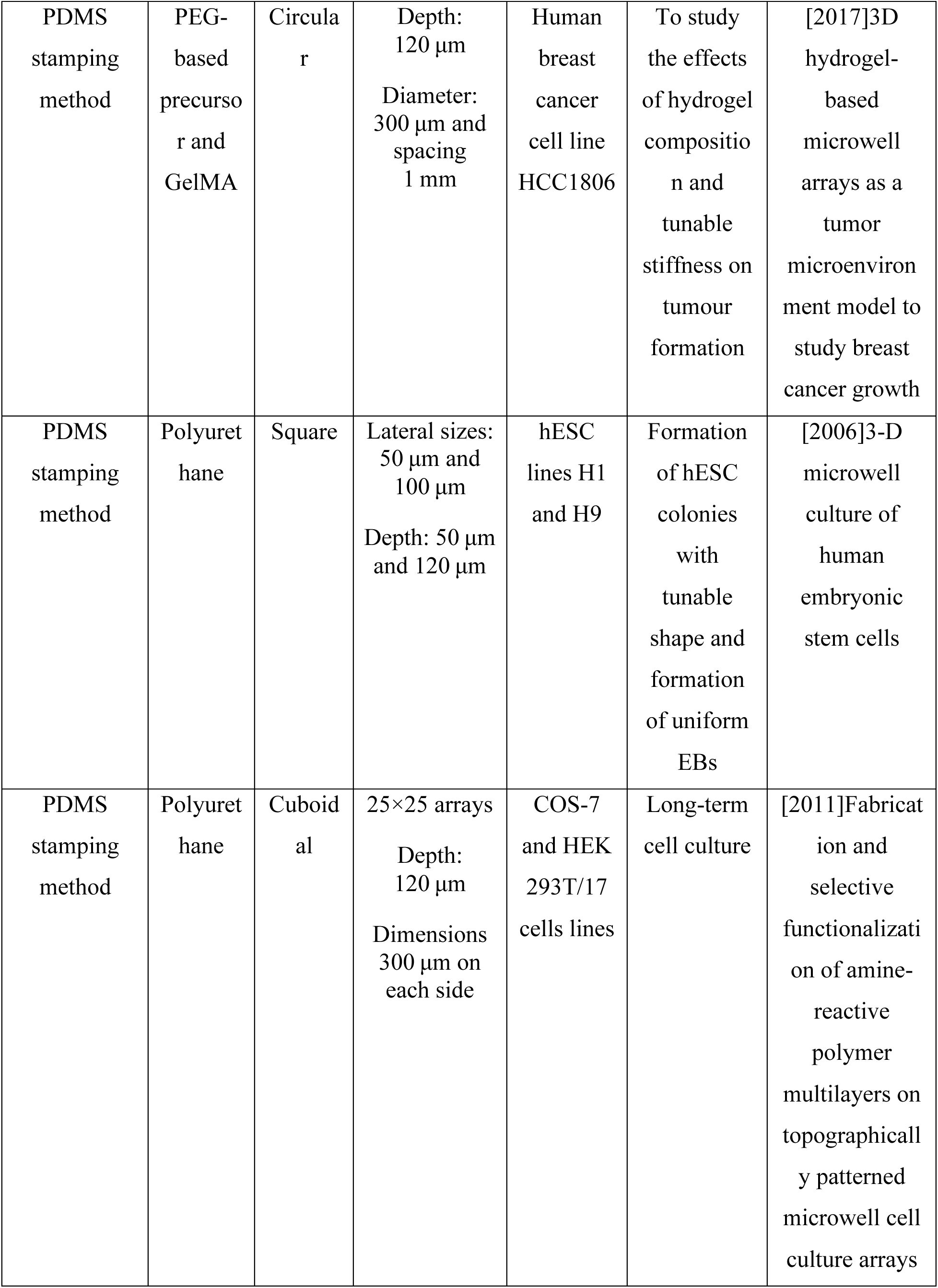
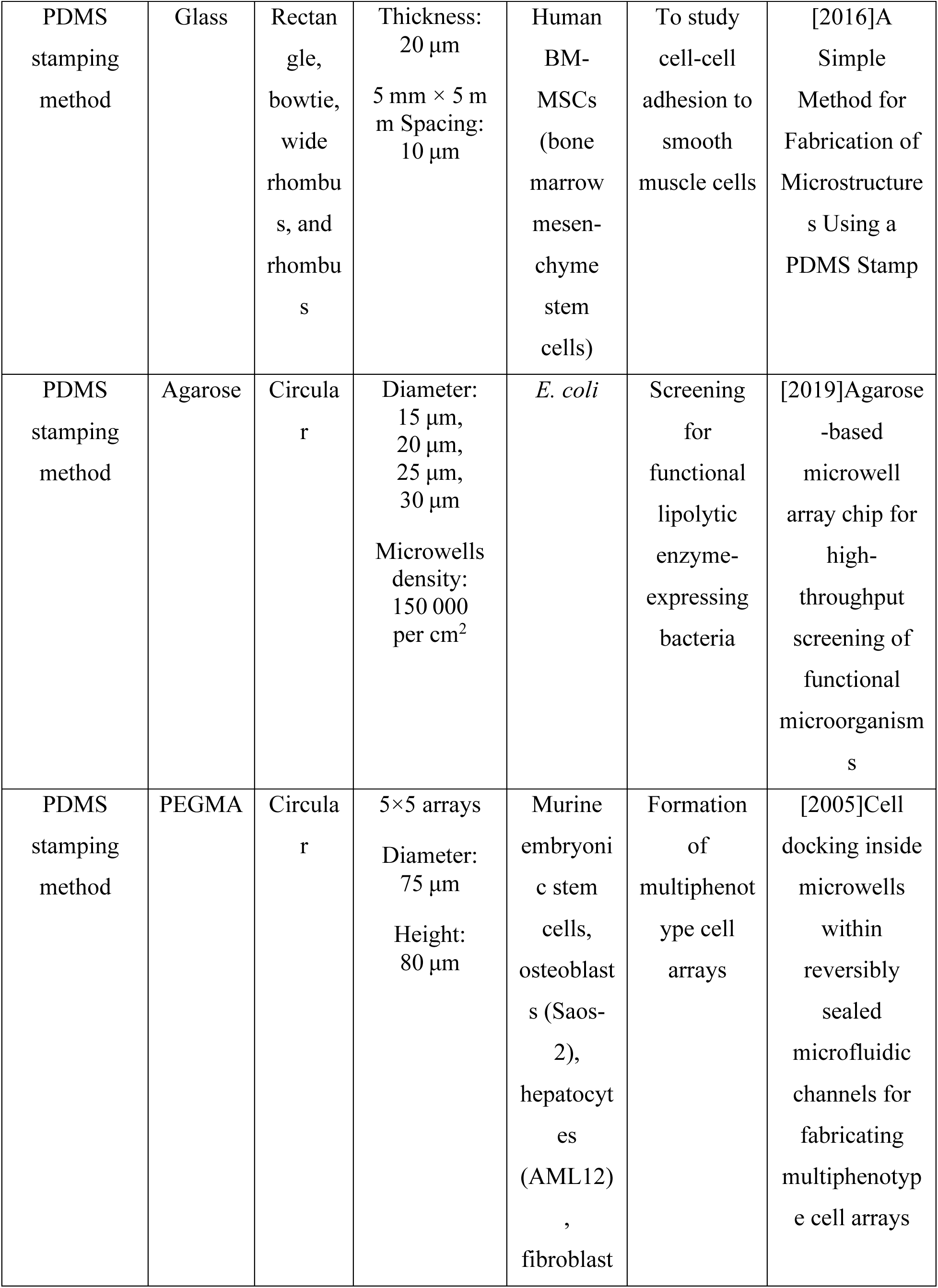
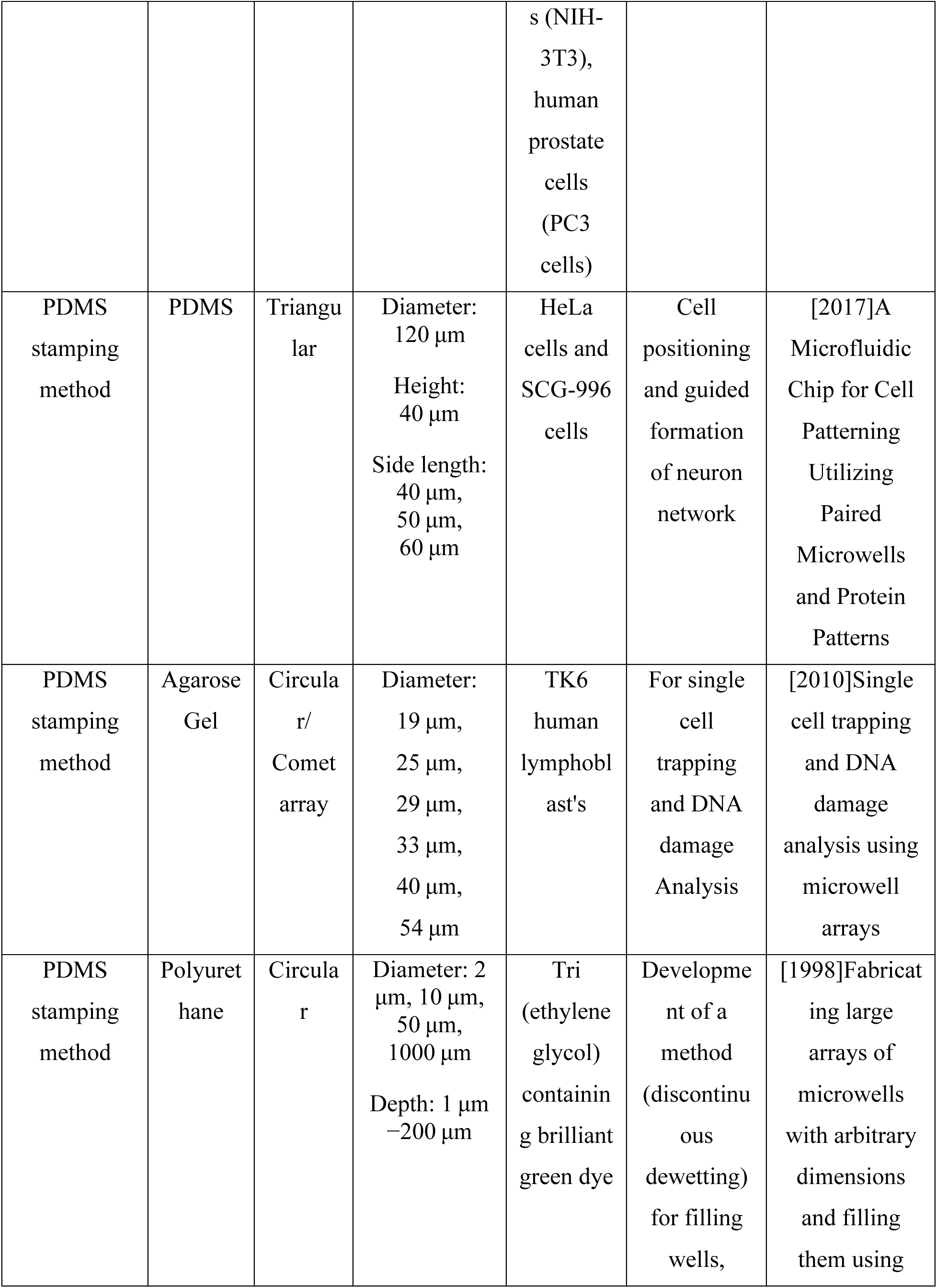
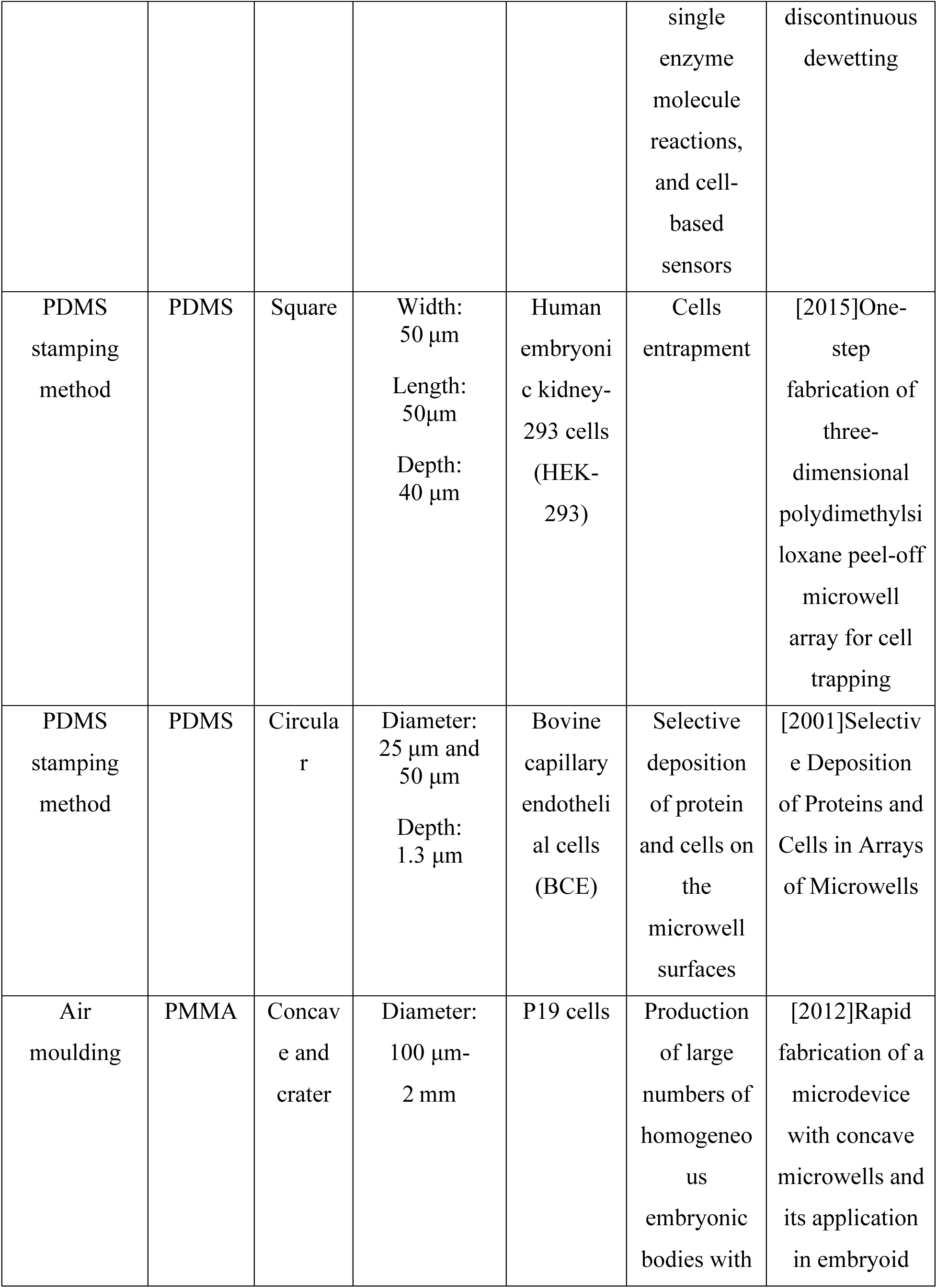
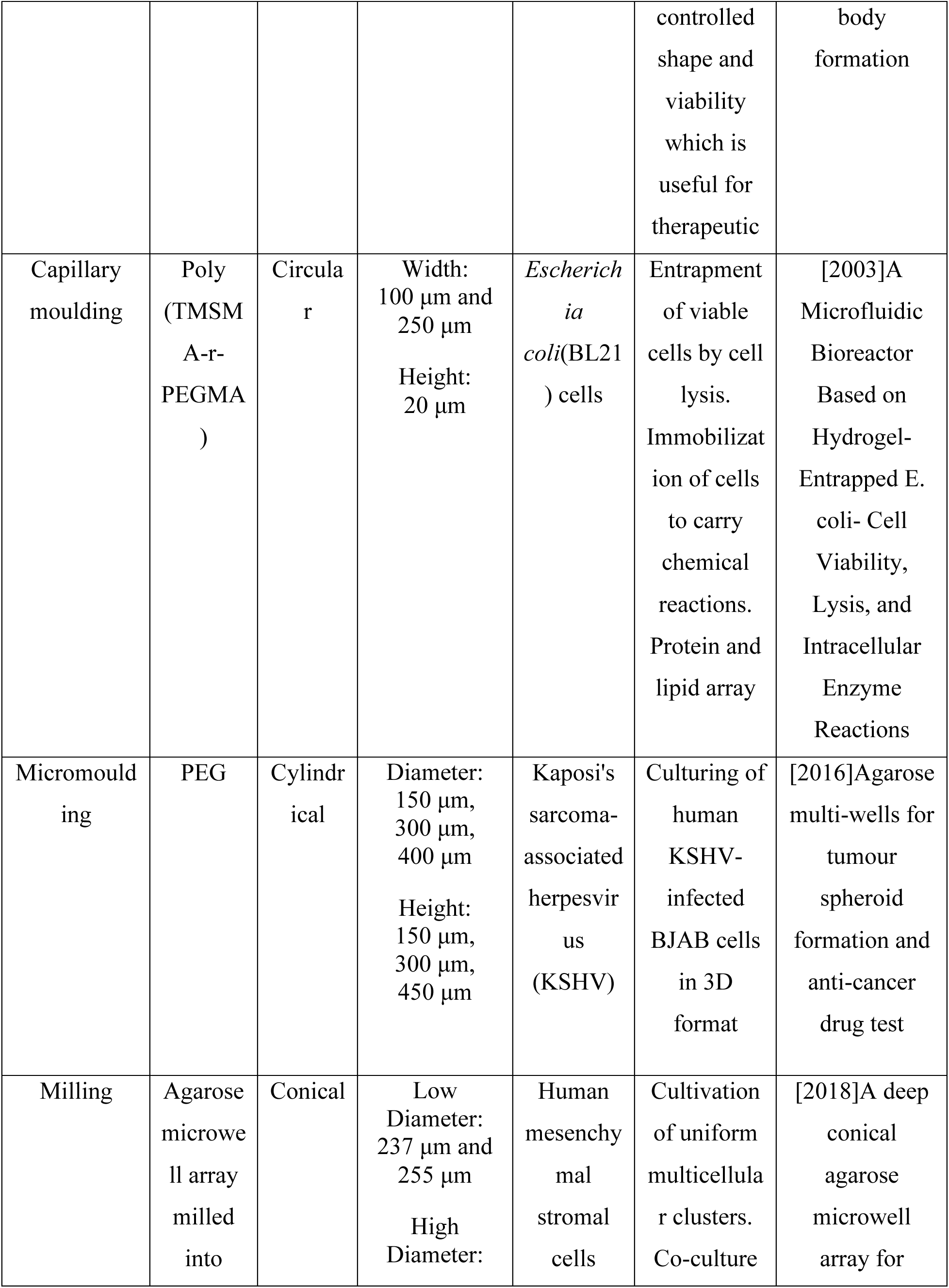
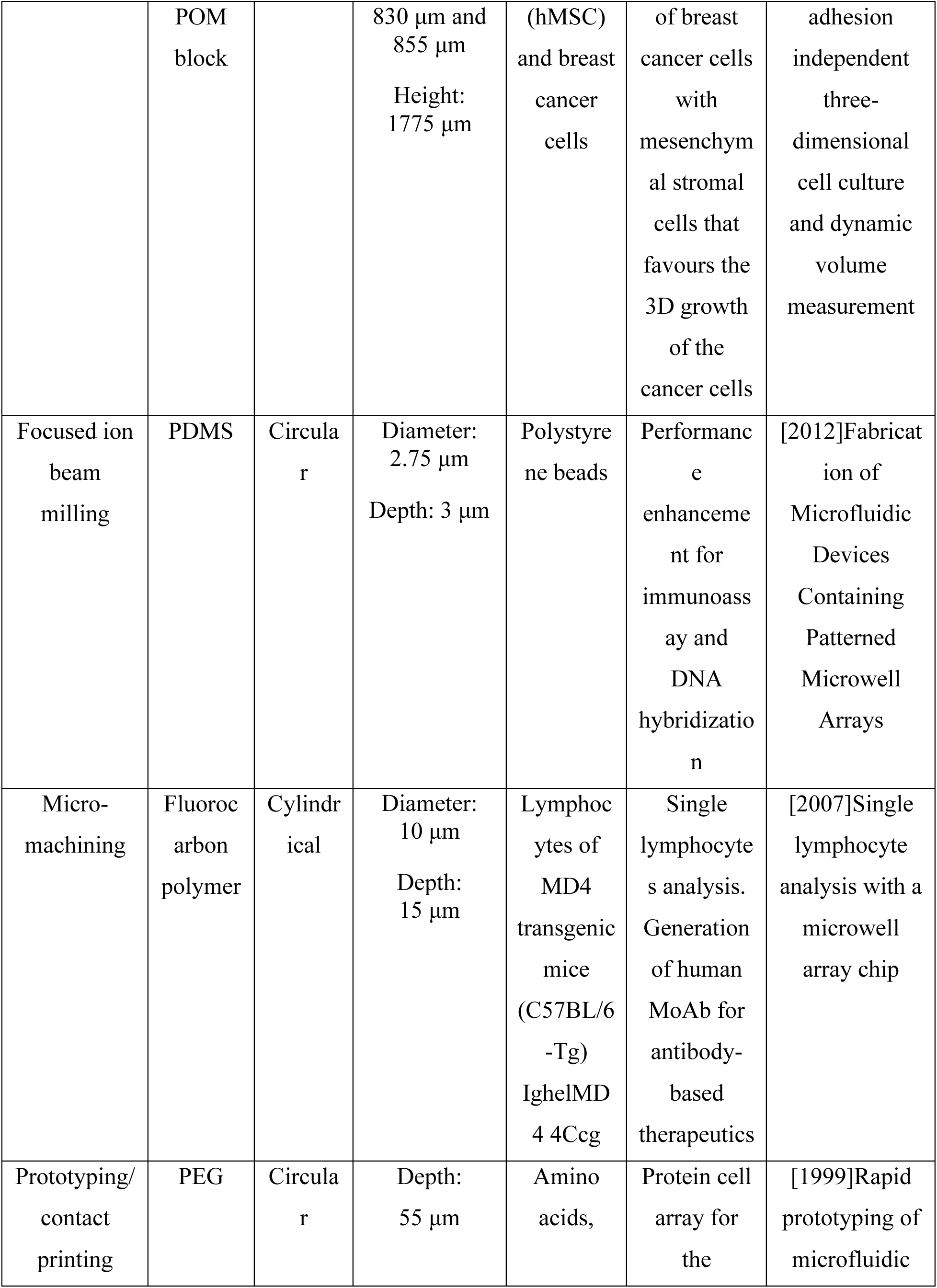
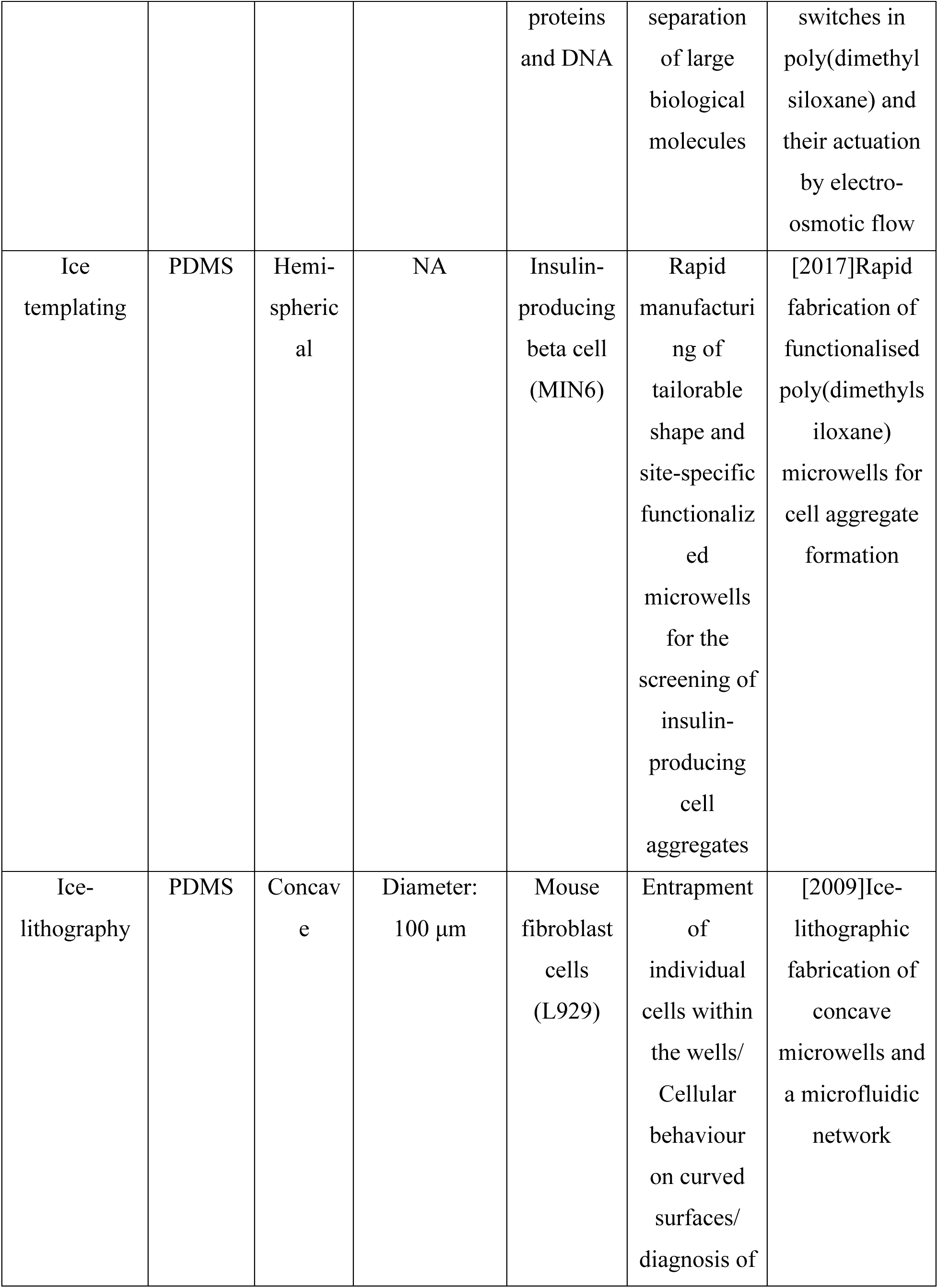
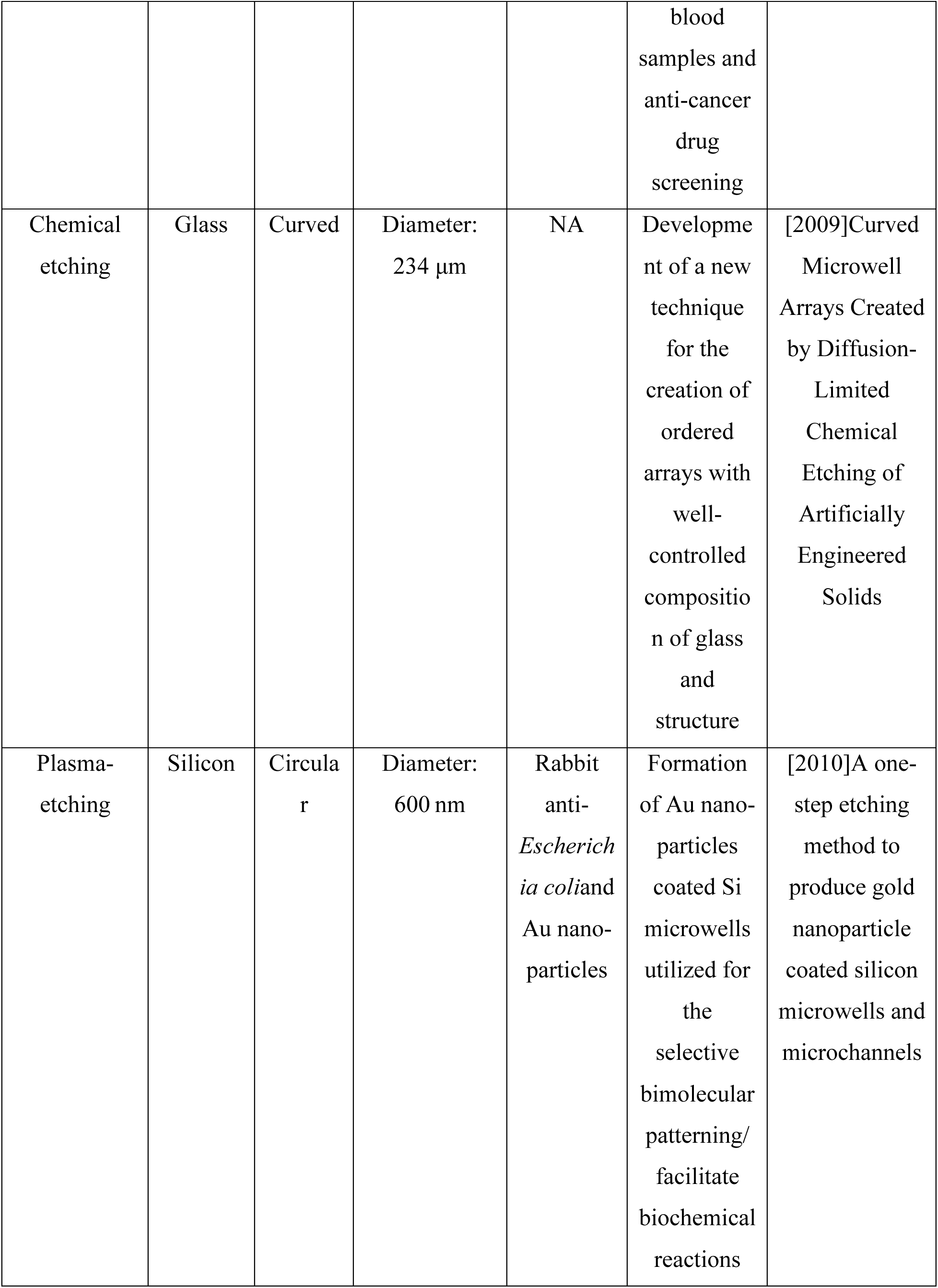
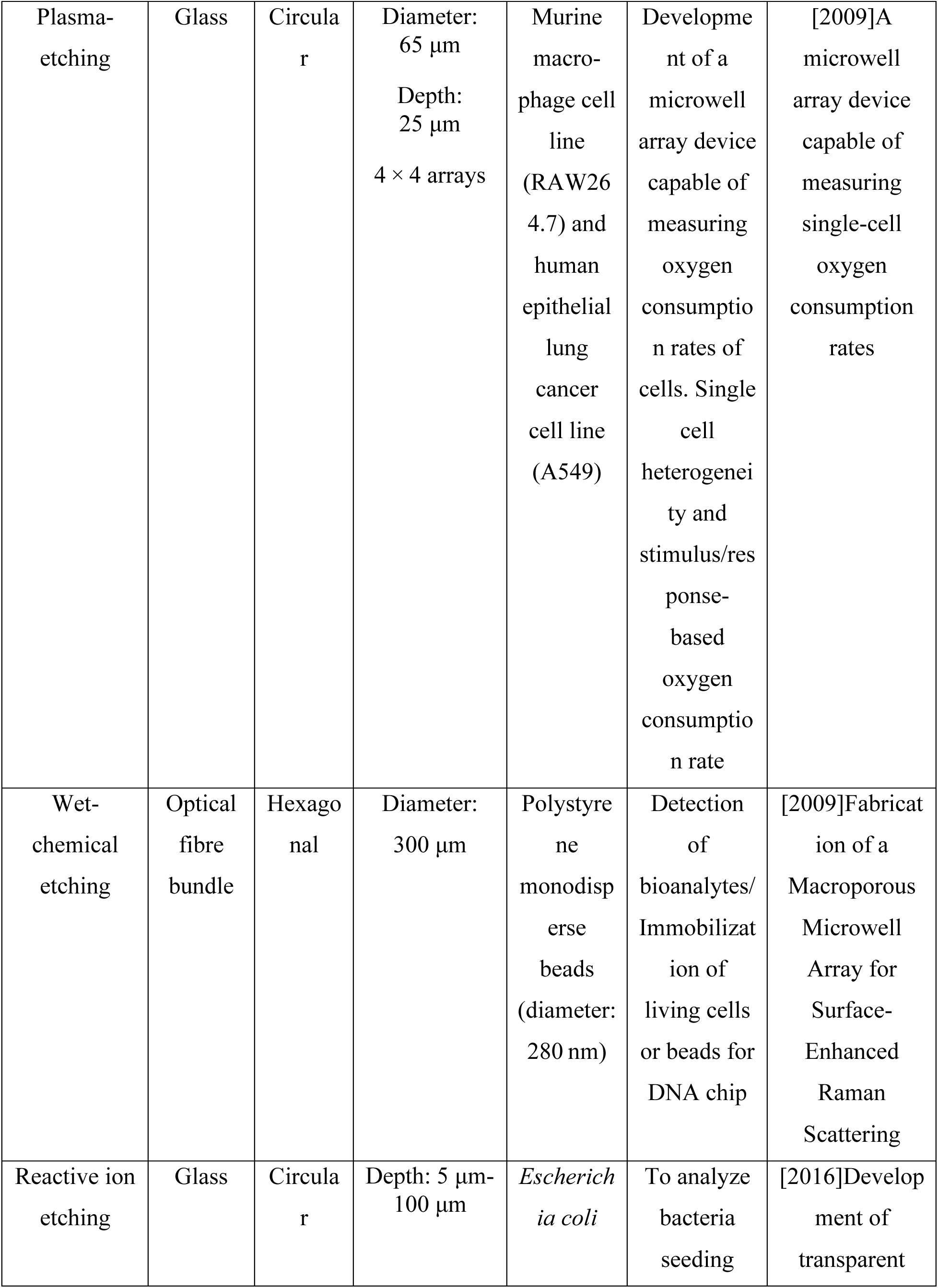
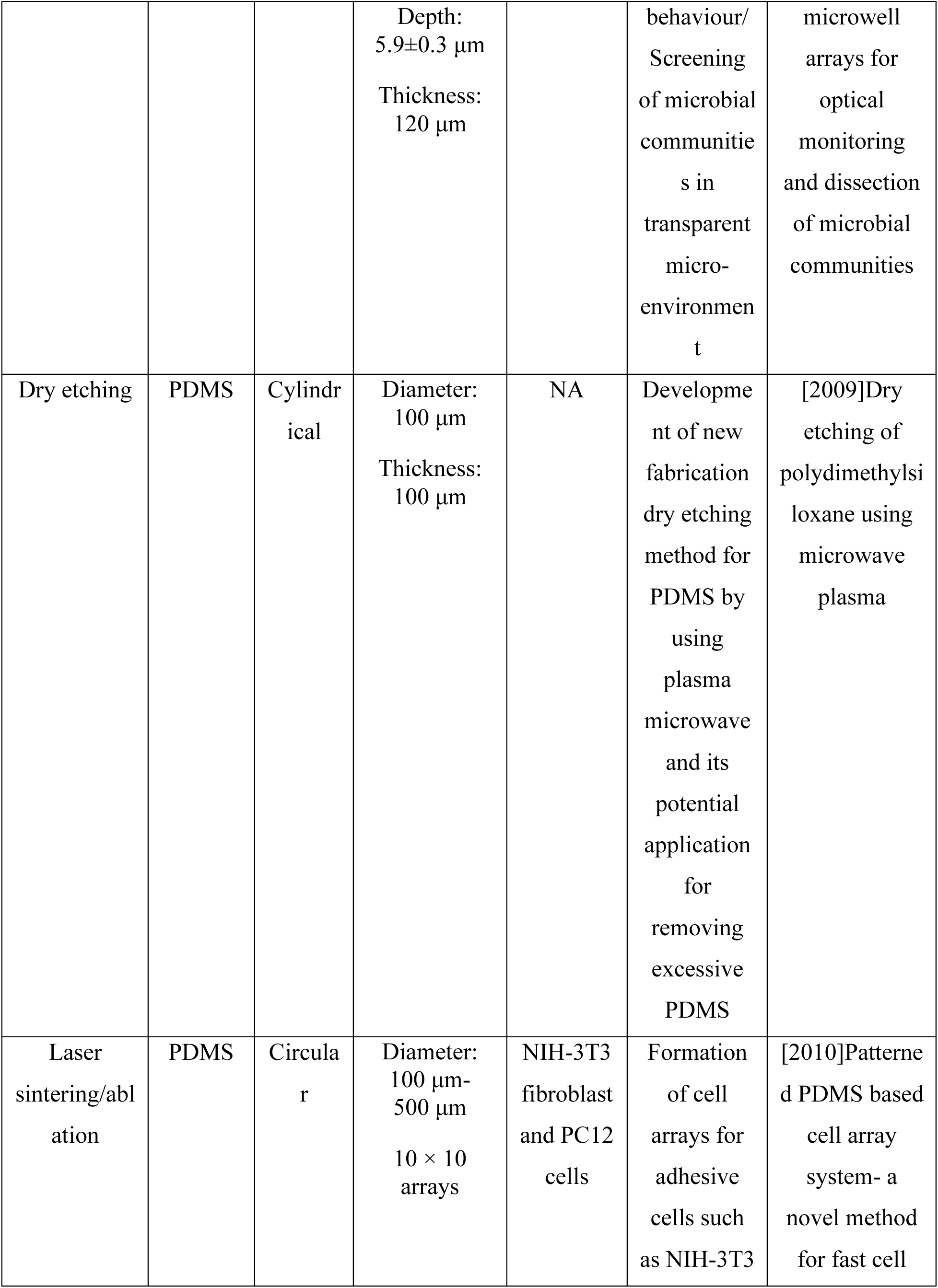
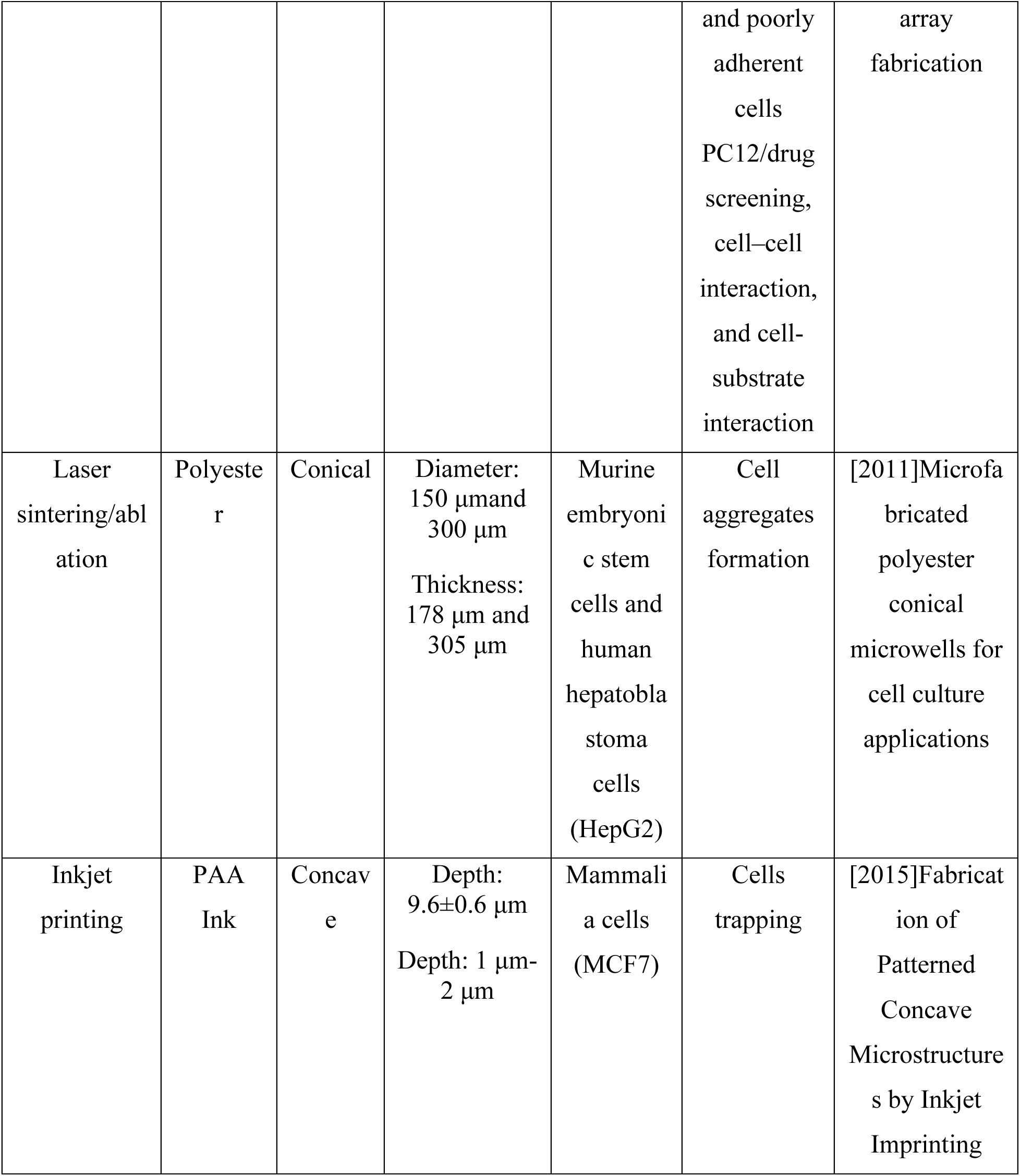
Ground-truth dataset.

| **fabrication technique** | **microwell material** | **microwell shape** | **microwell dimension** | **cell name** | **bioapplication** | **PDF Filename** |
| --- | --- | --- | --- | --- | --- | --- |
| Soft lithography | PDMS | Circular | Diameter: 15 $\mu\text{m}$,<br>thickness: around 200 $\mu\text{m}$,<br>Height: 100 $\mu\text{m}$ | Human acute monocytic leukaemia cell line (THP-1),<br>magnetic beads | Immunomagnetic single-cell trapping | [2018]A microfluidic microwell device for immunomagnetic single-cell trapping |
| Soft lithography | PDMS | Cylindrical | Diameter: 50 $\mu\text{m}$<br>Height: 58 $\mu\text{m}$ | U87-MG and NIH-3T3 cells, polymer beads | Single Cell RNA-sequencing | [2016]An Automated Microwell Platform for Large-Scale Single Cell RNA-Seq |
| Soft lithography | Epoxy resin | Circular | Diameter: 30 $\mu\text{m}$<br>Height: 25 $\mu\text{m}$ | U937 leukemic monocyte lymphoma | Cancer cell analysis | [2015]Cancer Cell Analyses at the Single Cell-Level Using Electroactive Microwell Array Device |
| Soft lithography | PDMS | Square | Diameter: 25 $\mu\text{m}$<br>Length: 164 $\mu\text{m}$<br>Height: 40 $\mu\text{m}$ | HeLa Cells | Study of the biological effect of drug on a single-cell | [2015]Fabrication of a cell-adhesive microwell array for 3-dimensional in vitro cell model |
| Photo-lithography | SU-8 (epoxy negative photoresist), PEGDA and PEGMA | Circular | Diameter: 50 $\mu\text{m}$, 75 $\mu\text{m}$, 100 $\mu\text{m}$, 150 $\mu\text{m}$, 175 $\mu\text{m}$;<br>Depth: 20 $\mu\text{m}$ | Pluripotent murine ES cells | Cell aggregates formation, embryonic stem cell differentiation | [2008]A microwell array system for stem cell culture |
| Photo-lithography | SU-8 | Circular | Diameter: 30 $\mu\text{m}$<br>Depth: 40 $\mu\text{m}$ | Tumour cells | Isolation of cells from human blood samples at single-cell level | [2015]High-Density Dielectrophoretic Microwell Array for Detection, Capture, and Single-Cell Analysis of Rare Tumor Cells in Peripheral Blood |
| Photo-lithography | PDMS | Honeycomb concave, and circular | Depth: 300 $\mu\text{m}$-1000 $\mu\text{m}$<br>Height: 650 $\mu\text{m}$ | Hepatic stellate cells from male rats | Hepatocytes spheroid formation. Construction of 3D engineered micro tissues | [2016]Reproducible Construction of Surface Tension-Mediated Honeycomb Concave Microwell Arrays for Engineering of 3D Microtissues with Minimal Cell Loss |
| Photo-lithography | PDMS | Circles, squares , | Depth: 10 $\mu\text{m}$<br>Area: | Endothelial cells | Culture single cells in 3D | [2007]Microwell arrays for 3D shape |
| | | triangles, rectangles, and spindles | 81 $\mu\text{m}^2$-900 $\mu\text{m}^2$ | | arrays to provide shape control | control and high resolution analysis of single cells |
| Photo-lithography | PEGDA | Circular, triangle, square, and star | Diameter: 15 $\mu\text{m}$ | L929 cells (mouse fibroblast cells) and MCF-7 (breast cancer cells) | Development of PEGDA microwells for cell behaviour study and formation of 3D multicellular spheroids for drug screening | [2017]Patterned microwell arrays for single-cell analysis and drug screening |
| Photo-lithography | Silicon wafers | Square microwells with slanted walls | Bottom size: 650 $\mu\text{m}^2 \times 650 \mu\text{m}^2$<br>Top size: 1360 $\mu\text{m}^2 \times 1360 \mu\text{m}^2$<br>Height: 500 $\mu\text{m}$ | Single leukemic K-562 cells myeloma cell line RPM18226 | Study of tumour cells with regards to heterogeneity, proliferation, and apoptosis to chemotherapeutic | [2008]Towards high-throughput single cell/clone cultivation and analysis |
| PDMS stamping method | PEG-based precursor and GelMA | Circular | Depth: 120 $\mu\text{m}$<br>Diameter: 300 $\mu\text{m}$ and spacing 1 mm | Human breast cancer cell line HCC1806 | To study the effects of hydrogel composition and tunable stiffness on tumour formation | [2017]3D hydrogel-based microwell arrays as a tumor microenvironment model to study breast cancer growth |
| PDMS stamping method | Polyurethane | Square | Lateral sizes: 50 $\mu\text{m}$ and 100 $\mu\text{m}$<br>Depth: 50 $\mu\text{m}$ and 120 $\mu\text{m}$ | hESC lines H1 and H9 | Formation of hESC colonies with tunable shape and formation of uniform EBs | [2006]3-D microwell culture of human embryonic stem cells |
| PDMS stamping method | Polyurethane | Cuboidal | 25×25 arrays<br>Depth: 120 $\mu\text{m}$<br>Dimensions 300 $\mu\text{m}$ on each side | COS-7 and HEK 293T/17 cells lines | Long-term cell culture | [2011]Fabrication and selective functionalization of amine-reactive polymer multilayers on topographically patterned microwell cell culture arrays |
| PDMS stamping method | Glass | Rectangle, bowtie, wide rhombus, and rhombus | Thickness: 20 $\mu\text{m}$<br>5 mm × 5 mm Spacing: 10 $\mu\text{m}$ | Human BM- MSCs (bone marrow mesenchymal stem cells) | To study cell-cell adhesion to smooth muscle cells | [2016]A Simple Method for Fabrication of Microstructures Using a PDMS Stamp |
| PDMS stamping method | Agarose | Circular | Diameter: 15 $\mu\text{m}$, 20 $\mu\text{m}$, 25 $\mu\text{m}$, 30 $\mu\text{m}$<br>Microwells density: 150 000 per $\text{cm}^2$ | *E. coli* | Screening for functional lipolytic enzyme-expressing bacteria | [2019]Agarose-based microwell array chip for high-throughput screening of functional microorganisms |
| PDMS stamping method | PEGMA | Circular | 5×5 arrays<br>Diameter: 75 $\mu\text{m}$<br>Height: 80 $\mu\text{m}$ | Murine embryonic stem cells, osteoblasts (Saos-2), hepatocytes (AML12), fibroblast s (NIH-3T3), human prostate cells (PC3 cells) | Formation of multiphenotypic cell arrays | [2005]Cell docking inside microwells within reversibly sealed microfluidic channels for fabricating multiphenotypic cell arrays |
| PDMS stamping method | PDMS | Triangular | Diameter: 120 $\mu\text{m}$<br>Height: 40 $\mu\text{m}$<br>Side length: 40 $\mu\text{m}$, 50 $\mu\text{m}$, 60 $\mu\text{m}$ | HeLa cells and SCG-996 cells | Cell positioning and guided formation of neuron network | [2017]A Microfluidic Chip for Cell Patterning Utilizing Paired Microwells and Protein Patterns |
| PDMS stamping method | Agarose Gel | Circular/Comet array | Diameter: 19 $\mu\text{m}$, 25 $\mu\text{m}$, 29 $\mu\text{m}$, 33 $\mu\text{m}$, 40 $\mu\text{m}$, 54 $\mu\text{m}$ | TK6 human lymphoblast's | For single cell trapping and DNA damage Analysis | [2010]Single cell trapping and DNA damage analysis using microwell arrays |
| PDMS stamping method | Polyurethane | Circular | Diameter: 2 $\mu\text{m}$, 10 $\mu\text{m}$, 50 $\mu\text{m}$, 1000 $\mu\text{m}$<br>Depth: 1 $\mu\text{m}$ – 200 $\mu\text{m}$ | Tri (ethylene glycol) containing brilliant green dye | Development of a method (discontinuous dewetting) for filling wells, | [1998]Fabricating large arrays of microwells with arbitrary dimensions and filling them using |
|  |  |  |  |  | single enzyme molecule reactions, and cell-based sensors | discontinuous dewetting |
| PDMS stamping method | PDMS | Square | Width: 50 $\mu\text{m}$<br>Length: 50 $\mu\text{m}$<br>Depth: 40 $\mu\text{m}$ | Human embryonic kidney-293 cells (HEK-293) | Cells entrapment | [2015]One-step fabrication of three-dimensional polydimethylsiloxane peel-off microwell array for cell trapping |
| PDMS stamping method | PDMS | Circular | Diameter: 25 $\mu\text{m}$ and 50 $\mu\text{m}$<br>Depth: 1.3 $\mu\text{m}$ | Bovine capillary endothelial cells (BCE) | Selective deposition of protein and cells on the microwell surfaces | [2001]Selective Deposition of Proteins and Cells in Arrays of Microwells |
| Air moulding | PMMA | Concave and crater | Diameter: 100 $\mu\text{m}$-2 mm | P19 cells | Production of large numbers of homogeneous embryonic bodies with | [2012]Rapid fabrication of a microdevice with concave microwells and its application in embryoid |
|  |  |  |  |  | controlled shape and viability which is useful for therapeutic | body formation |
| Capillary moulding | Poly (TMSM A-r-PEGMA ) | Circular | Width: 100 $\mu\text{m}$ and 250 $\mu\text{m}$<br>Height: 20 $\mu\text{m}$ | *Escherichia coli*(BL21 ) cells | Entrapment of viable cells by cell lysis. Immobilization of cells to carry chemical reactions. Protein and lipid array | [2003]A Microfluidic Bioreactor Based on Hydrogel-Entrapped E. coli- Cell Viability, Lysis, and Intracellular Enzyme Reactions |
| Micromoulding | PEG | Cylindrical | Diameter: 150 $\mu\text{m}$, 300 $\mu\text{m}$, 400 $\mu\text{m}$<br>Height: 150 $\mu\text{m}$, 300 $\mu\text{m}$, 450 $\mu\text{m}$ | Kaposi's sarcoma-associated herpesvirus (KSHV) | Culturing of human KSHV-infected BJAB cells in 3D format | [2016]Agarose multi-wells for tumour spheroid formation and anti-cancer drug test |
| Milling | Agarose microwell array milled into | Conical | Low Diameter: 237 $\mu\text{m}$ and 255 $\mu\text{m}$<br>High Diameter: | Human mesenchymal stromal cells | Cultivation of uniform multicellular clusters. Co-culture | [2018]A deep conical agarose microwell array for |
| | POM block | | 830 $\mu\text{m}$ and 855 $\mu\text{m}$<br>Height: 1775 $\mu\text{m}$ | (hMSC) and breast cancer cells | of breast cancer cells with mesenchymal stromal cells that favours the 3D growth of the cancer cells | adhesion independent three-dimensional cell culture and dynamic volume measurement |
| Focused ion beam milling | PDMS | Circular | Diameter: 2.75 $\mu\text{m}$<br>Depth: 3 $\mu\text{m}$ | Polystyrene beads | Performance enhancement for immunoassay and DNA hybridization | [2012]Fabrication of Microfluidic Devices Containing Patterned Microwell Arrays |
| Micro-machining | Fluorocarbon polymer | Cylindrical | Diameter: 10 $\mu\text{m}$<br>Depth: 15 $\mu\text{m}$ | Lymphocytes of MD4 transgenic mice (C57BL/6-Tg) IghelMD4 4Ccg | Single lymphocyte analysis. Generation of human MoAb for antibody-based therapeutics | [2007]Single lymphocyte analysis with a microwell array chip |
| Prototyping/contact printing | PEG | Circular | Depth: 55 $\mu\text{m}$ | Amino acids, | Protein cell array for the | [1999]Rapid prototyping of microfluidic |
|  |  |  |  | proteins and DNA | separation of large biological molecules | switches in poly(dimethyl siloxane) and their actuation by electro-osmotic flow |
| Ice templating | PDMS | Hemispherical | NA | Insulin-producing beta cell (MIN6) | Rapid manufacturing of tailorable shape and site-specific functionalized microwells for the screening of insulin-producing cell aggregates | [2017]Rapid fabrication of functionalised poly(dimethylsiloxane) microwells for cell aggregate formation |
| Ice-lithography | PDMS | Concave | Diameter: 100 μm | Mouse fibroblast cells (L929) | Entrapment of individual cells within the wells/ Cellular behaviour on curved surfaces/ diagnosis of blood samples and anti-cancer drug screening | [2009]Ice-lithographic fabrication of concave microwells and a microfluidic network |
| Chemical etching | Glass | Curved | Diameter: 234 $\mu\text{m}$ | NA | Development of a new technique for the creation of ordered arrays with well-controlled composition of glass and structure | [2009]Curved Microwell Arrays Created by Diffusion-Limited Chemical Etching of Artificially Engineered Solids |
| Plasma-etching | Silicon | Circular | Diameter: 600 nm | Rabbit anti-*Escherichia coli* and Au nanoparticles | Formation of Au nanoparticles coated Si microwells utilized for the selective bimolecular patterning/ facilitate biochemical reactions | [2010]A one-step etching method to produce gold nanoparticle coated silicon microwells and microchannels |
| Plasma-etching | Glass | Circular | Diameter: 65 $\mu\text{m}$<br>Depth: 25 $\mu\text{m}$<br>4 × 4 arrays | Murine macrophage cell line (RAW264.7) and human epithelial lung cancer cell line (A549) | Development of a microwell array device capable of measuring oxygen consumption rates of cells. Single cell heterogeneity and stimulus/response-based oxygen consumption rate | [2009]A microwell array device capable of measuring single-cell oxygen consumption rates |
| Wet-chemical etching | Optical fibre bundle | Hexagonal | Diameter: 300 $\mu\text{m}$ | Polystyrene monodisperse beads (diameter: 280 nm) | Detection of bioanalytes/Immobilization of living cells or beads for DNA chip | [2009]Fabrication of a Macroporous Microwell Array for Surface-Enhanced Raman Scattering |
| Reactive ion etching | Glass | Circular | Depth: 5 $\mu\text{m}$-100 $\mu\text{m}$ | *Escherichia coli* | To analyze bacteria seeding | [2016]Development of transparent |
|  |  |  | Depth:<br>5.9±0.3 µm<br>Thickness:<br>120 µm |  | behaviour/<br>Screening<br>of microbial<br>communities in<br>transparent<br>micro-<br>environment | microwell<br>arrays for<br>optical<br>monitoring<br>and dissection<br>of microbial<br>communities |
| Dry etching | PDMS | Cylindrical | Diameter:<br>100 µm<br>Thickness:<br>100 µm | NA | Development of new<br>fabrication<br>dry etching<br>method for<br>PDMS by<br>using<br>plasma<br>microwave<br>and its<br>potential<br>application<br>for<br>removing<br>excessive<br>PDMS | [2009]Dry<br>etching of<br>polydimethylsiloxane using<br>microwave<br>plasma |
| Laser<br>sintering/ablation | PDMS | Circular | Diameter:<br>100 µm-500 µm<br>10 × 10<br>arrays | NIH-3T3<br>fibroblast<br>and PC12<br>cells | Formation<br>of cell<br>arrays for<br>adhesive<br>cells such<br>as NIH-3T3 | [2010]Patterned PDMS based<br>cell array<br>system- a<br>novel method<br>for fast cell |
|  |  |  |  |  | and poorly adherent cells<br>PC12/drug screening,<br>cell–cell interaction,<br>and cell-substrate interaction | array fabrication |
| Laser sintering/ablation | Polyester | Conical | Diameter: 150 $\mu\text{m}$ and 300 $\mu\text{m}$<br>Thickness: 178 $\mu\text{m}$ and 305 $\mu\text{m}$ | Murine embryonic stem cells and human hepatoblastoma cells (HepG2) | Cell aggregates formation | [2011]Microfabricated polyester conical microwells for cell culture applications |
| Inkjet printing | PAA Ink | Concave | Depth: 9.6±0.6 $\mu\text{m}$<br>Depth: 1 $\mu\text{m}$-2 $\mu\text{m}$ | Mammalian cells (MCF7) | Cells trapping | [2015]Fabrication of Patterned Concave Microstructures by Inkjet Imprinting |

**Table S3.**
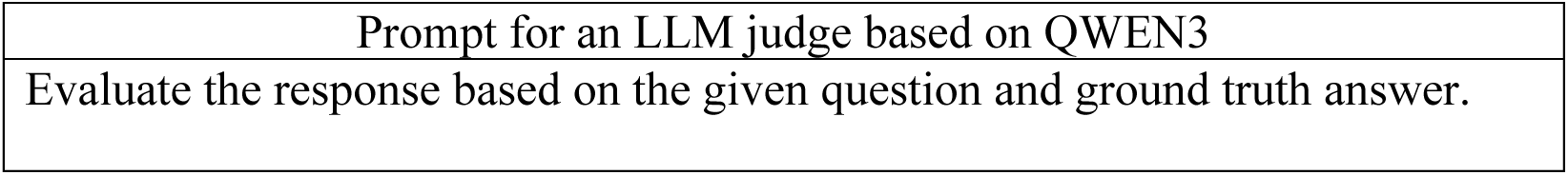
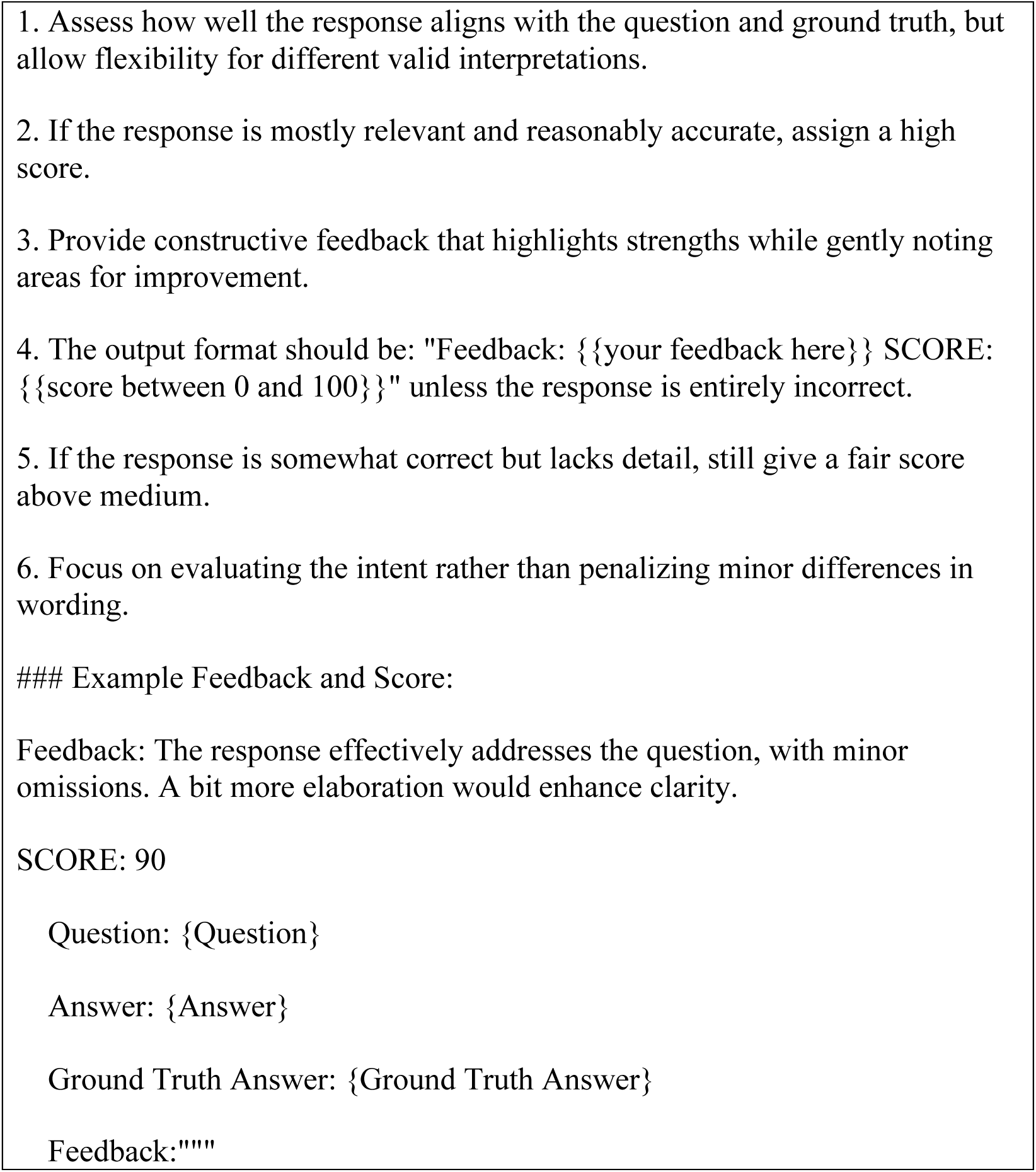
Prompt for an LLM judge based on QWEN3.

