## Supplementary Information for "MicrowellMicrofluidicsMiner (M³): Leverage Large Language Model Agents for Knowledge Mining of Microwell Microfluidics"

*Corresponding Author

*Supplementary* *File*

QWEN3-extractor’s Prompt

Context: {Context}
Question: {Question}
Responses from LLMs:

LLM1 response: {LLM1 response}
LLM2 response: {LLM2 response}
LLM3 response: {LLM3 response}
LLM4 response: {LLM4 response}
You are an expert judge evaluating responses from four different large language models (LLMs) to a given question.
Your task is to select the best response strictly from the provided options based on the retrieved context, following a structured decision-making process.
<INSTRUCTION>
1. Choose exactly one response from the four provided. You must not modify or generate a new response.
2. Evaluate responses based on their accuracy, clarity, completeness, and coherence with both the question and the retrieved context.
3. Follow the selection process below:
Selection Process:
Step 1: Majority Voting Rule
If two or more LLMs provide the same response, select that response as the final answer.
Step 2: Tie-Breaking Rule (if no majority exists)
If there is no majority but two responses are equally strong, choose the response that is:
Most factually supported by the retrieved context.
More concise and well-structured while preserving accuracy.
Step 3: Fallback Mechanism (if all four responses differ)
If all four responses are unique, select the most contextually relevant and factually accurate response according to the retrieved context.
If multiple responses are equally strong, apply the tie-breaking rule from Step 2.
Return only the selected response, without any additional explanation or commentary.
<INSTRUCTION>
<FORMAT>
Answer: Return only the selected response, without any additional explanation or commentary.
<FORMAT>

Table S1. QWEN3-extractor’s Prompt

| **fabrication technique** | **microwell material** | **microwell shape** | **microwell dimension** | **cell name** | **bioapplication** | **PDF Filename** |
| --- | --- | --- | --- | --- | --- | --- |
| Soft lithography | PDMS | Circular | Diameter: 15 μm, thickness: around 200 μm, Height: 100 μm | Human acute monocytic leukaemia cell line (THP-1), magnetic beads | Immunomagnetic single-cell trapping | [2018]A microfluidic microwell device for immunomagnetic single-cell trapping |
| Soft lithography | PDMS | Cylindrical | Diameter: 50 μm  Height: 58 μm | U87-MG and NIH-3T3 cells, polymer beads | Single Cell RNA-sequencing | [2016]An Automated Microwell Platform for Large-Scale Single Cell RNA-Seq |
| Soft lithography | Epoxy resin | Circular | Diameter: 30 μm  Height: 25 μm | U937 leukemic monocyte lymphoma | Cancer cell analysis | [2015]Cancer Cell Analyses at the Single Cell-Level Using Electroactive Microwell Array Device |
| Soft lithography | PDMS | Square | Diameter: 25 μm  Length: 164 μm  Height: 40 μm | HeLa Cells | Study of the biological effect of drug on a single-cell | [2015]Fabrication of a cell-adhesive microwell array for 3-dimensional in vitro cell model |
| Photo-lithography | SU-8 (epoxy negative photoresist), PEGDA and PEGMA | Circular | Diameter: 50 μm, 75 μm, 100 μm, 150 μm, 175 μm; Depth: 20 μm | Pluri-potent murine ES cells | Cell aggregates formation, embryonic stem cell differentiation | [2008]A microwell array system for stem cell culture |
| Photo-lithography | SU-8 | Circular | Diameter: 30 μm  Depth: 40 μm | Tumour cells | Isolation of cells from human blood samples at single-cell level | [2015]High-Density Dielectrophoretic Microwell Array for Detection, Capture, and Single-Cell Analysis of Rare Tumor Cells in Peripheral Blood |
| Photo-lithography | PDMS | Honeycomb concave, and circular | Depth: 300 μm-1000 μm  Height: 650 μm | Hepatic stellate cells from male rats | Hepatocytes spheroid formation. Construction of 3D engineered micro tissues | [2016]Reproducible Construction of Surface Tension-Mediated Honeycomb Concave Microwell Arrays for Engineering of 3D Microtissues with Minimal Cell Loss |
| Photo-lithography | PDMS | Circles, squares, triangles, rectangles, and spindles | Depth: 10 μm Area: 81 μm^2^-900 μm^2^ | Endothelial cells | Culture single cells in 3D arrays to provide shape control | [2007]Micro-well arrays for 3D shape control and high resolution analysis of single cells |
| Photo-lithography | PEGDA | Circular, triangle, square, and star | Diameter: 15 μm | L929 cells (mouse fibroblast cells) and MCF-7 (breast cancer cells) | Development of PEGDA microwells for cell behaviour study and formation of 3D multicellular spheroids for drug screening | [2017]Patterned microwell arrays for single-cell analysis and drug screening |
| Photo-lithography | Silicon wafers | Square microwells with slanted walls | Bottom size: 650 μm^2^×650 μm^2^  Top size: 1360 μm^2^×1360 μm^2^  Height: 500 μm | Single leukemic K-562 cells myeloma cell line RPM18226 | Study tumour cells with regards to heterogeneity, proliferation, and apoptosis to chemotherapeutic | [2008]Towards high-throughput single cell/clone cultivation and analysis |
| PDMS stamping method | PEG-based precursor and GelMA | Circular | Depth: 120 μm  Diameter: 300 μm and spacing 1 mm | Human breast cancer cell line HCC1806 | To study the effects of hydrogel composition and tunable stiffness on tumour formation | [2017]3D hydrogel-based microwell arrays as a tumor microenvironment model to study breast cancer growth |
| PDMS stamping method | Polyurethane | Square | Lateral sizes: 50 μm and 100 μm  Depth: 50 μm and 120 μm | hESC lines H1 and H9 | Formation of hESC colonies with tunable shape and formation of uniform EBs | [2006]3-D microwell culture of human embryonic stem cells |
| PDMS stamping method | Polyurethane | Cuboidal | 25×25 arrays  Depth: 120 μm  Dimensions 300 μm on each side | COS-7 and HEK 293T/17 cells lines | Long-term cell culture | [2011]Fabrication and selective functionalization of amine-reactive polymer multilayers on topographically patterned microwell cell culture arrays |
| PDMS stamping method | Glass | Rectangle, bowtie, wide rhombus, and rhombus | Thickness: 20 μm  5 mm × 5 mm Spacing: 10 μm | Human BM-MSCs (bone marrow mesen-chyme stem cells) | To study cell-cell adhesion to smooth muscle cells | [2016]A Simple Method for Fabrication of Microstructures Using a PDMS Stamp |
| PDMS stamping method | Agarose | Circular | Diameter: 15 μm, 20 μm, 25 μm, 30 μm  Microwells density: 150 000 per cm^2^ | *E. coli* | Screening for functional lipolytic enzyme-expressing bacteria | [2019]Agarose-based microwell array chip for high-throughput screening of functional microorganisms |
| PDMS stamping method | PEGMA | Circular | 5×5 arrays  Diameter: 75 μm  Height: 80 μm | Murine embryonic stem cells, osteoblasts (Saos-2), hepatocytes (AML12), fibroblasts (NIH-3T3), human prostate cells (PC3 cells) | Formation of multiphenotype cell arrays | [2005]Cell docking inside microwells within reversibly sealed microfluidic channels for fabricating multiphenotype cell arrays |
| PDMS stamping method | PDMS | Triangular | Diameter: 120 μm  Height: 40 μm  Side length: 40 μm, 50 μm, 60 μm | HeLa cells and SCG-996 cells | Cell positioning and guided formation of neuron network | [2017]A Microfluidic Chip for Cell Patterning Utilizing Paired Microwells and Protein Patterns |
| PDMS stamping method | Agarose Gel | Circular/ Comet array | Diameter: 19 μm, 25 μm, 29 μm, 33 μm, 40 μm, 54 μm | TK6 human lymphoblast's | For single cell trapping and DNA damage Analysis | [2010]Single cell trapping and DNA damage analysis using microwell arrays |
| PDMS stamping method | Polyurethane | Circular | Diameter: 2 μm, 10 μm, 50 μm, 1000 μm  Depth: 1 μm −200 μm | Tri (ethylene glycol) containing brilliant green dye | Development of a method (discontinuous dewetting) for filling wells, single enzyme molecule reactions, and cell-based sensors | [1998]Fabricating large arrays of microwells with arbitrary dimensions and filling them using discontinuous dewetting |
| PDMS stamping method | PDMS | Square | Width: 50 μm  Length: 50μm  Depth: 40 μm | Human embryonic kidney-293 cells (HEK-293) | Cells entrapment | [2015]One-step fabrication of three-dimensional polydimethylsiloxane peel-off microwell array for cell trapping |
| PDMS stamping method | PDMS | Circular | Diameter: 25 μm and 50 μm  Depth: 1.3 μm | Bovine capillary endothelial cells (BCE) | Selective deposition of protein and cells on the microwell surfaces | [2001]Selective Deposition of Proteins and Cells in Arrays of Microwells |
| Air moulding | PMMA | Concave and crater | Diameter: 100 μm-2 mm | P19 cells | Production of large numbers of homogeneous embryonic bodies with controlled shape and viability which is useful for therapeutic | [2012]Rapid fabrication of a microdevice with concave microwells and its application in embryoid body formation |
| Capillary moulding | Poly (TMSMA-r-PEGMA) | Circular | Width: 100 μm and 250 μm  Height: 20 μm | *Escherichia coli*(BL21) cells | Entrapment of viable cells by cell lysis. Immobilization of cells to carry chemical reactions. Protein and lipid array | [2003]A Microfluidic Bioreactor Based on Hydrogel-Entrapped E. coli- Cell Viability, Lysis, and Intracellular Enzyme Reactions |
| Micromoulding | PEG | Cylindrical | Diameter: 150 μm, 300 μm, 400 μm  Height: 150 μm, 300 μm, 450 μm | Kaposi's sarcoma-associated herpesvirus (KSHV) | Culturing of human KSHV-infected BJAB cells in 3D format | [2016]Agarose multi-wells for tumour spheroid formation and anti-cancer drug test |
| Milling | Agarose microwell array milled into POM block | Conical | Low Diameter: 237 μm and 255 μm  High Diameter: 830 μm and 855 μm  Height: 1775 μm | Human mesenchymal stromal cells (hMSC) and breast cancer cells | Cultivation of uniform multicellular clusters. Co-culture of breast cancer cells with mesenchymal stromal cells that favours the 3D growth of the cancer cells | [2018]A deep conical agarose microwell array for adhesion independent three-dimensional cell culture and dynamic volume measurement |
| Focused ion beam milling | PDMS | Circular | Diameter: 2.75 μm  Depth: 3 μm | Polystyrene beads | Performance enhancement for immunoassay and DNA hybridization | [2012]Fabrication of Microfluidic Devices Containing Patterned Microwell Arrays |
| Micro-machining | Fluorocarbon polymer | Cylindrical | Diameter: 10 μm  Depth: 15 μm | Lymphocytes of MD4 transgenic mice (C57BL/6-Tg) IghelMD4 4Ccg | Single lymphocytes analysis. Generation of human MoAb for antibody-based therapeutics | [2007]Single lymphocyte analysis with a microwell array chip |
| Prototyping/contact printing | PEG | Circular | Depth: 55 μm | Amino acids, proteins and DNA | Protein cell array for the separation of large biological molecules | [1999]Rapid prototyping of microfluidic switches in poly(dimethyl siloxane) and their actuation by electro-osmotic flow |
| Ice templating | PDMS | Hemi-spherical | NA | Insulin-producing beta cell (MIN6) | Rapid manufacturing of tailorable shape and site-specific functionalized microwells for the screening of insulin-producing cell aggregates | [2017]Rapid fabrication of functionalised poly(dimethylsiloxane) microwells for cell aggregate formation |
| Ice-lithography | PDMS | Concave | Diameter: 100 μm | Mouse fibroblast cells (L929) | Entrapment of individual cells within the wells/ Cellular behaviour on curved surfaces/ diagnosis of blood samples and anti-cancer drug screening | [2009]Ice-lithographic fabrication of concave microwells and a microfluidic network |
| Chemical etching | Glass | Curved | Diameter: 234 μm | NA | Development of a new technique for the creation of ordered arrays with well-controlled composition of glass and structure | [2009]Curved Microwell Arrays Created by Diffusion-Limited Chemical Etching of Artificially Engineered Solids |
| Plasma-etching | Silicon | Circular | Diameter: 600 nm | Rabbit anti-*Escherichia coli*and Au nano-particles | Formation of Au nano-particles coated Si microwells utilized for the selective bimolecular patterning/ facilitate biochemical reactions | [2010]A one-step etching method to produce gold nanoparticle coated silicon microwells and microchannels |
| Plasma-etching | Glass | Circular | Diameter: 65 μm  Depth: 25 μm  4 × 4 arrays | Murine macro-phage cell line (RAW264.7) and human epithelial lung cancer cell line (A549) | Development of a microwell array device capable of measuring oxygen consumption rates of cells. Single cell heterogeneity and stimulus/response-based oxygen consumption rate | [2009]A microwell array device capable of measuring single-cell oxygen consumption rates |
| Wet-chemical etching | Optical fibre bundle | Hexagonal | Diameter: 300 μm | Polystyrene monodisperse beads (diameter: 280 nm) | Detection of bioanalytes/ Immobilization of living cells or beads for DNA chip | [2009]Fabrication of a Macroporous Microwell Array for Surface-Enhanced Raman Scattering |
| Reactive ion etching | Glass | Circular | Depth: 5 μm-100 μm  Depth: 5.9±0.3 μm  Thickness: 120 μm | *Escherichia coli* | To analyze bacteria seeding behaviour/ Screening of microbial communities in transparent micro-environment | [2016]Development of transparent microwell arrays for optical monitoring and dissection of microbial communities |
| Dry etching | PDMS | Cylindrical | Diameter: 100 μm  Thickness: 100 μm | NA | Development of new fabrication dry etching method for PDMS by using plasma microwave and its potential application for removing excessive PDMS | [2009]Dry etching of polydimethylsiloxane using microwave plasma |
| Laser sintering/ablation | PDMS | Circular | Diameter: 100 μm-500 μm  10 × 10 arrays | NIH-3T3 fibroblast and PC12 cells | Formation of cell arrays for adhesive cells such as NIH-3T3 and poorly adherent cells PC12/drug screening, cell–cell interaction, and cell-substrate interaction | [2010]Patterned PDMS based cell array system- a novel method for fast cell array fabrication |
| Laser sintering/ablation | Polyester | Conical | Diameter: 150 μmand 300 μm  Thickness: 178 μm and 305 μm | Murine embryonic stem cells and human hepatoblastoma cells (HepG2) | Cell aggregates formation | [2011]Microfabricated polyester conical microwells for cell culture applications |
| Inkjet printing | PAA Ink | Concave | Depth: 9.6±0.6 μm  Depth: 1 μm-2 μm | Mammalia cells (MCF7) | Cells trapping | [2015]Fabrication of Patterned Concave Microstructures by Inkjet Imprinting |

Table S2. Ground-truth dataset

Prompt for an LLM judge based on QWEN3

Evaluate the response based on the given question and ground truth answer.

1. Assess how well the response aligns with the question and ground truth, but allow flexibility for different valid interpretations.

2. If the response is mostly relevant and reasonably accurate, assign a high score.

3. Provide constructive feedback that highlights strengths while gently noting areas for improvement.

4. The output format should be: "Feedback: {{your feedback here}} SCORE: {{score between 0 and 100}}" unless the response is entirely incorrect.

5. If the response is somewhat correct but lacks detail, still give a fair score above medium.

6. Focus on evaluating the intent rather than penalizing minor differences in wording.

\### Example Feedback and Score:

Feedback: The response effectively addresses the question, with minor omissions. A bit more elaboration would enhance clarity.

SCORE: 90

Question: {Question}

Answer: {Answer}

Ground Truth Answer: {Ground Truth Answer}

Feedback:"""

Table S3. Prompt for an LLM judge based on QWEN3
